# Rab40/Cullin5 complex regulates EPLIN and actin cytoskeleton dynamics during cell migration and invasion

**DOI:** 10.1101/2021.04.01.438077

**Authors:** Erik S Linklater, Emily D Duncan, Ke-Jun Han, Algirdas Kaupinis, Mindaugas Valius, Traci R Lyons, Rytis Prekeris

## Abstract

Rab40b is a SOCS box containing protein that regulates the secretion of MMPs to facilitate extracellular matrix remodeling during cell migration. Here we show that Rab40b interacts with Cullin5 via the Rab40b SOCS domain. We demonstrate that loss of Rab40b/Cullin5 binding decreases cell motility and invasive potential, and show that defective cell migration and invasion stem from alteration to the actin cytoskeleton, leading to decreased invadopodia formation, decreased actin dynamics at the leading edge, and an increase in stress fibers. We also show that these stress fibers anchor at less dynamic, more stable focal adhesions. Mechanistically, changes in the cytoskeleton and focal adhesion dynamics are mediated in part by EPLIN, which we demonstrate to be a binding partner of Rab40b and a target for Rab40b/Cullin5 dependent localized ubiquitylation and degradation. Thus, we propose a model where the Rab40b/Cullin5 dependent ubiquitylation regulates EPLIN localization to promote cell migration and invasion by altering focal adhesion and cytoskeletal dynamics.

## INTRODUCTION

Cell migration is a complex and highly regulated process that involves coordinated changes in signaling, membrane trafficking and cytoskeleton dynamics. Consequently, during epithelial-to-mesenchymal-transition (EMT) in development or carcinogenesis, cells undergo an extensive shift in genetic and post-translational programming to promote cellular pathways important for migration, such as the loss of cell-cell adhesion and enhanced localized dynamics of both the actin cytoskeleton and focal adhesions (FAs) [1, 2]. Additionally, it is well-established that extracellular matrix (ECM) degradation and remodeling plays a key role in mediating cell migration during development and cancer metastasis. ECM remodeling is facilitated via the delivery and secretion of matrix metalloproteinases (MMPs) to the leading edge of invasive cells [3–7].

While the cellular machinery mediating this targeted release remains to be fully defined, it is clear that MMPs are released at cellular extensions known as migratory pseudopods or podosomes in normal cells, or invadopodia in cancer cells [8, 9]. Regardless of the context, these cellular extensions are typically formed and extended via localized polymerization of the actin cytoskeleton and occur with the coordinated assembly/disassembly of focal adhesion sites (FAs) [10–12]. Thus, the key to cell migration through the ECM is the coordination between actin polymerization, FA assembly/disassembly and targeted secretion of MMPs. How all these processes are integrated and regulated during cell migration remains to be fully understood and is a main focus of this study.

Since MMP targeting to invadopodia is one of the key events during cancer cell migration, we and others aim to identify regulators of this process. Rab GTPases are known master regulators of targeted vesicle transport and cargo secretion [13]. Accordingly, roles for various Rabs in the delivery of MT1-MMP (also known as MMP14), including Rab5a, Rab8, Rab14, and Rab27a have been established [14–16]. Work from our lab specifically identified Rab40b as an important regulator of MMP2/9 secretion and ECM remodeling during breast cancer cell invasion. Relatedly, we have also demonstrated that Tks5, a known invadopodia regulatory protein, is an effector protein for Rab40b [17, 18]. Additional work from other labs has further corroborated an emerging role for Rab40b in cell migration and cancer progression [19–22].

The Rab40 sub-family contains four closely related isoforms: Rab40a, Rab40al, Rab40b and Rab40c, with Rab40a and Rab40al being expressed only in simian primates. This sub-family is unique amongst the Rab GTPases due to the presence of a C-Terminal SOCS box motif (Figure 1A) [23], which allows binding to members of the Cullin family of proteins [24, 25]. Cullins are the central component of the largest class of E3 ubiquitin ligases, referred to as Cullin-RING ligases (CRLs) [26, 27].

**Figure 1.**
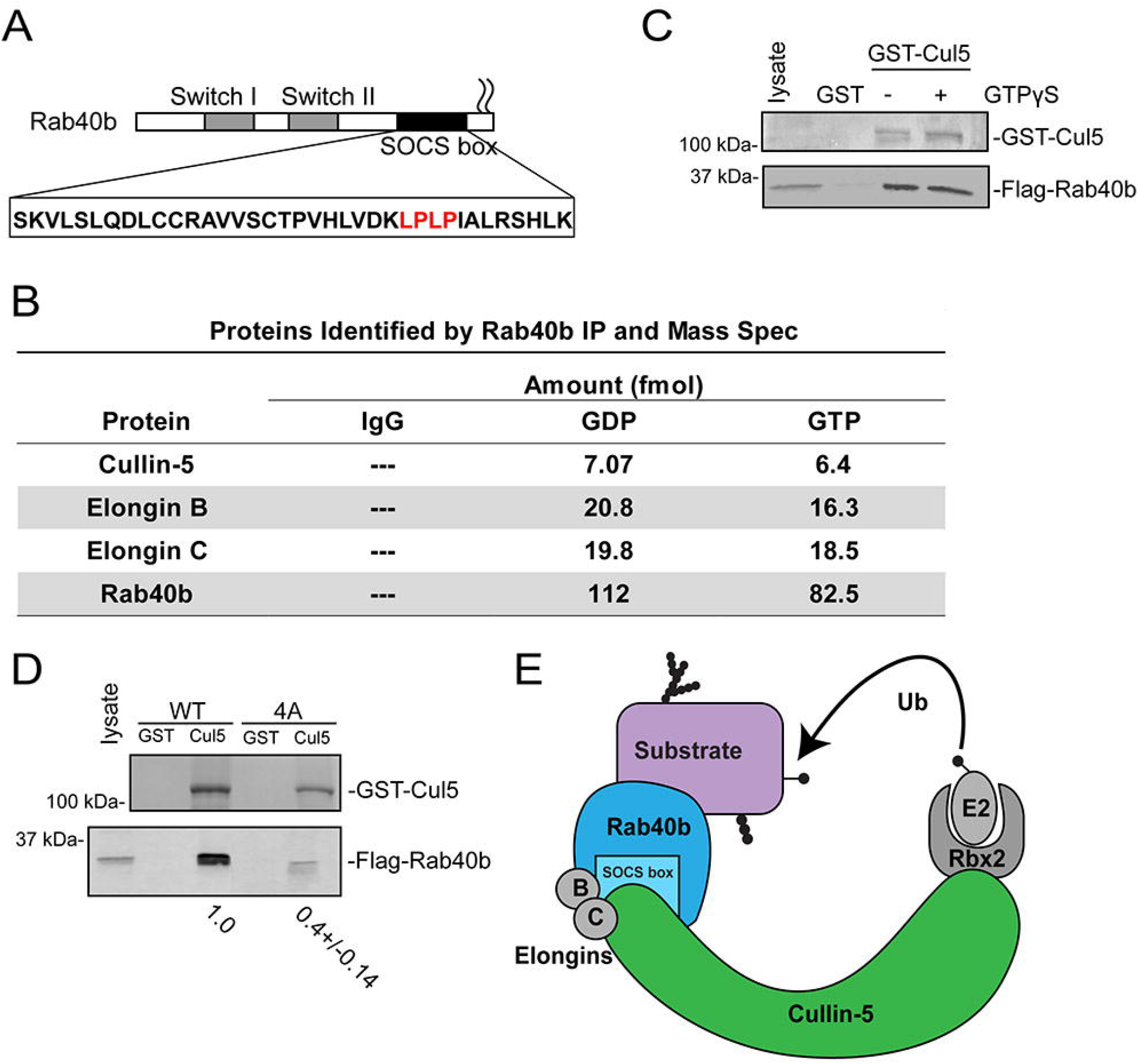
Rab40b binds to Cullin5 in SOCS-dependent and GTP independent manner. (A) Schematic diagram of Rab40b. (B) List of proteins identified in both GDP and GTPγS bound Flag-Rab40b immunoprecipitate from MDA-MB-231 lysate followed by mass spectroscopy analysis. (C) Flag-Rab40b binding to GST-Cullin 5 as analyzed by glutathione bead pull-down assay. Top panel, coomassie staining. Bottom panel, Western Blot with anti-Flag. (D) Flag-Rab40b (wild type and 4A mutants) binding to GST-Cullin5 as analyzed by glutathione bead pulldown assay followed by Western Blot with anti-Flag (bottom blot) and coomassie stain (top blot). Numbers shown are the average densitometry analysis derived from three independent experiments, standardized to wild type Flag-Rab40b. Wild type::4A, p < 0.001. (F) Model showing CRL5 E3 ligase complex partners.

Within the Cullin5 specific CRL (CRL5), Cullin5 acts as a scaffold protein that recruits RING-box protein Rbx2, the Elongin B/C complex, and a SOCS box-containing adaptor protein. Together, this CRL5 complex mediates ubiquitylation of SOCS-bound substrate molecules. Cullin/SOCS complexes, first identified as scaffolds for JAK/STAT signaling [28–30] regulate a wide range of substrate proteins, including VHL [31], DAB1 [32], and p130Cas [32]. The canonical SOCS-containing family of eight proteins has steadily been expanded to over forty [33], including the Rab40 sub-family. Both Rab40a and Rab40c complex with Cullin5 to ubiquitylate Pak4 [34] and Rack1 [35], respectively, ultimately resulting in their proteosomal degradation. However, little is known about the function of the Rab40b in this context, particularly in the setting of cell migration, invadopodia formation and ECM remodeling. Furthermore, it remains unclear how much functional redundancy exists among the Rab40 isoforms.

We hypothesized that the Rab40b SOCS box would allow interaction with the CRL5 complex and contribute to coordination of MMP secretion by inducing changes in actin and FA dynamics. Consistent with this hypothesis, we demonstrate that Rab40b binds to Cullin5 in a SOCS box dependent fashion and that formation of this CRL5 complex is required for chemotactic migration and breast cancer cell invasion. We also show this Rab40b/Cullin5 complex influences multiple aspects of cell migration, including FA localization and dynamics, stress fiber induction, formation of invadopodia, and changes in lamellipodia dynamics. In this study we completed an unbiased proteomics screen to identify Rab40b/Cullin5 specific substrate proteins. One of the proteins identified was EPLIN (Epithelial Protein Lost In Neoplasia, encoded by the LIMA1 gene), a tumor suppressor known to inhibit cell migration and EMT [36–40]. Here we demonstrate that preventing Rab40b/Cullin5 binding increases total cellular EPLIN levels in breast cancer cells and causes its re-distribution to stress fibers and the leading edge of migratory lamellipodia. Additionally, we show that increased cellular EPLIN contributes to changes in focal adhesion and actin dynamics. Finally, we demonstrate that ablation of Rab40b/Cullin5 complex formation inhibits tumor growth and EMT *in vivo* in a xenograft model.

## RESULTS

### Rab40b Binds to Cullin5 in SOCS-Dependent and GTP-Independent Manner

Identification of the full complement of Rab40b interacting partners is necessary to define how Rab40b contributes to both cell migration and other biological processes. To identify additional Rab40b binding partners we first performed a proteomics screen. Due to the lack of reliable commercial antibodies for Rab40b, we used a previously generated triple-negative breast cancer (TNBC) cell line MDA-MB-231 that stably expresses Flag–Rab40b [17]. Lysates from these cells were incubated with anti-Flag-antibody conjugated beads or control IgG beads. Because Rab GTPases cycle between inactive GDP-bound and active GTP-bound states, we also immunoprecipitated Flag-Rab40b in the presence of GDP or the non-hydrolysable GTP analog GTPγS. Bound proteins were then analyzed by mass spectrometry. Cullin5 as well as CRL5 complex partners Elongin B and Elongin C were identified as putative Rab40b interacting proteins (Figure 1B and Supplemental Figure 1A). All three proteins bound relatively equally to GDP or GTPγS bound Rab40b, suggesting that binding of the CRL5 complex is independent of the nucleotide state of Rab40b. To confirm that Cullin5 binds to Rab40b, we next incubated lysates from MDA-MB-231 cells constitutively expressing Flag-Rab40b with glutathione beads conjugated to GST-only or purified recombinant GST-Cullin5. Consistent with our mass spec results, Flag-Rab40b interacted with GST-Cullin5 independent of its nucleotide state (Figure 1C).

It is well established that a highly conserved LPLP motif, named the Cul-Box, is found within the SOCS box and that this motif is necessary for the binding of Cullin complexes to canonical SOCS proteins [33]. This motif is also present in Rab40b (Figure 1A, in red). To determine whether this motif is important for the binding of Rab40b to Cullin5 we mutated the LPLP sequence to AAAA (Flag-Rab40b-4A) which has been shown to abate the binding of Cullin5 to other SOCS-containing proteins [31]. Lysates from MDA-MB-231 cell lines stably expressing either wild-type Flag-Rab40b or Flag-Rab40b-4A were used to test the ability of Rab40b to bind to GST-Cullin5 using glutathione bead pull-down assays. As shown in Figure 1D, mutant Flag-Rab40b-4A has significantly reduced binding to GST-Cullin5 compared to wild-type. Taken together, these results demonstrate that mammalian Rab40b binds to Cullin5 and Elongin B/C independent of its nucleotide state, and that binding to Cullin5 is mediated by the Rab40b SOCS box (Figure 1E).

### Rab40b/Cullin5 Regulates Cell Migration via Cytoskeletal Reorganization

Given that Cullin family members and their substrates are important regulators of cell migration [41–44], together with an emerging role for Rab40b in cell motility, we hypothesized that Rab40b and Cullin5 binding may be an important regulator of this process. Using MDA-MB-231 cells stably expressing either wild-type Flag-Rab40b or the Cullin5 binding deficient Flag-Rab40b-4A mutant (Figure 2A), we first sought to assess potential differences in migration using scratch wound assay. Somewhat surprisingly, expression of either wildtype or mutant Rab40b-4A had no discernable effect on collective cell migration (Supplemental Figure 2A). Analysis of individual cells can often reveal changes in migration dynamics that can be missed using population migration assays, such as scratch wound assay. Thus, we sought to look for differences in individual cell movement. Control or Flag-Rab40b-4A-expressing cells were plated on collagen-coated dishes, placed in full medium, stained with SiR-actin and DAPI and imaged once every 10 minutes for up to 10 hours and individually tracked (Supplemental Movies 1 and 2). Generally, Flag-Rab40b-4A-expressing cells were capable of moving in similar fashion to control cells, suggesting that disrupting Rab40b and Cullin5 interaction does not inhibit the cells ability to move. Tracking of individual cells allowed for quantification of various properties of cell migration, such as velocity, directionality, and persistence. Mean Squared Displacement (MSD), which quantifies the randomness of movement by assessing the amount of space explored by a particle within a system, showed no significant difference between control and Flag-Rab40b-4A cells. When the MSD data is fit to the power-law function MSD(Δt) = C*Δt^α^, wherein the exponent α is indicative of the type of motion observed, both control and Rab40b-4A cells exhibit similar values that are representative of superdiffusive motion [45, 46] (Supplemental Figure 1B). Additionally, Flag-Rab40b-4A cells showed no significant difference in velocity compared to control MDA-MB-231 cells (Supplemental Figure 1C). Finally, we sought to assess “straightness” of cell movement by determining the cell directionality ratio, a measure of the net displacement of a cell from its starting to final position compared to the total distance traveled. Interestingly, Flag-Rab40b-4A-expressing cells exhibit a slight but significant decrease in directionality (Figure 2B, Supplemental Figure 1D), a phenotype that would be difficult to ascertain during the collective cell migration that underpins scratch wound assays.

**Figure 2.**
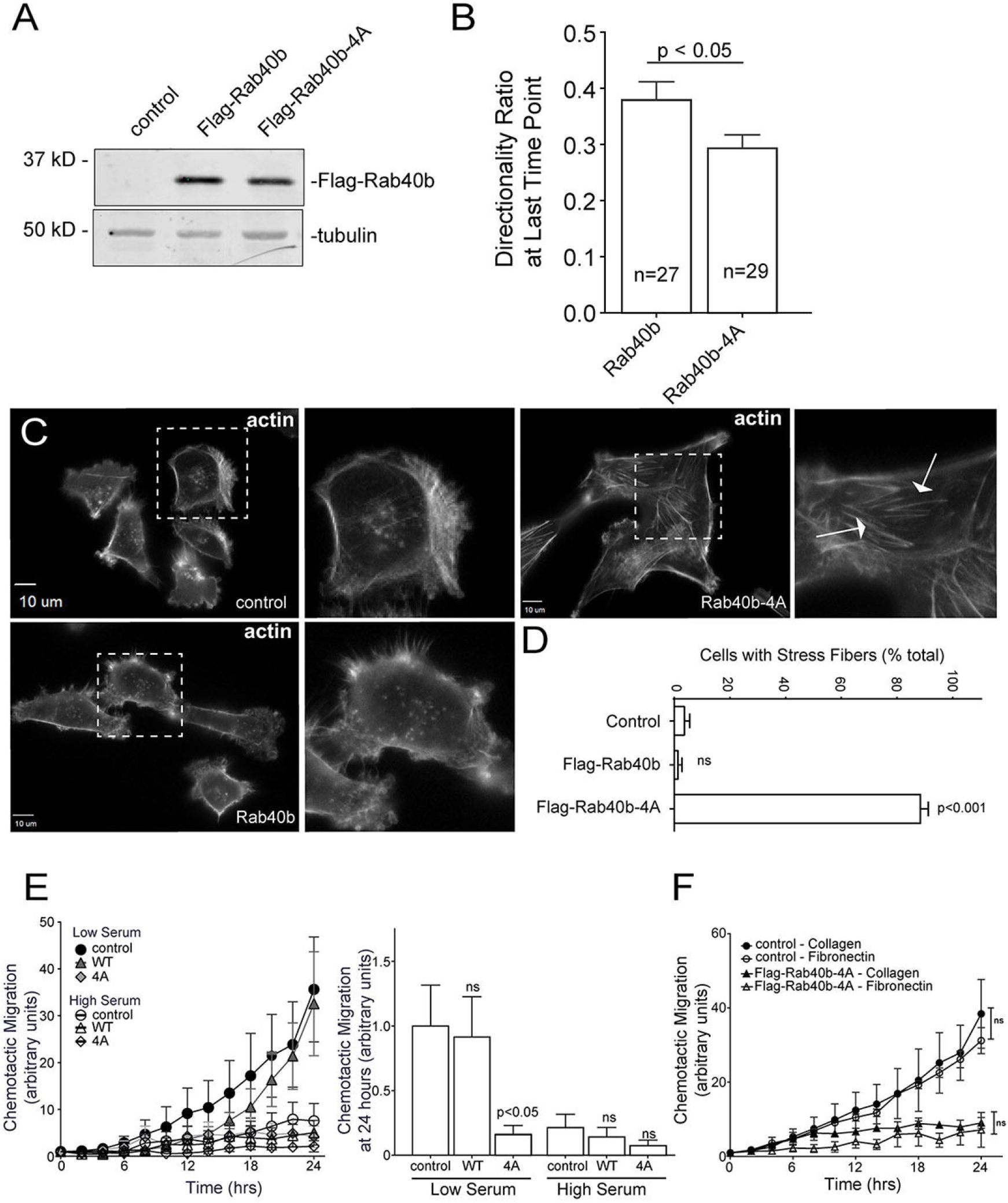
Rab40b/Cullin5 Affects Individual Cell Migration. (A) Western Bot analysis of lysates from MDA-MB-231 cells stably expressing wild type Flag-Rab40b or Flag-Rab40b-4A. (B) Directionality of migrating cells derived from time-lapse analysis of control or Flag-Rab40b-4A cells (see Supplemental Movies 1 and 2) for cells that remained in frame for the duration of the experiment. Shown data are the means and standard error of means. n is the number of cells analyzed. (C-D) Control, Flag-Rab40b, Flag-Rab40b-4A cells were plated on collagen-coated coverslips, fixed and stained with phalloidin-Alexa594. Zoomed regions of interest highlight differences in cytoskeletal architecture and arrows point to stress-fibers. (D) Quantification of cells with prominent stress fibers. n ≥ 100 cells per condition. (E) Chemotactic assays of control, Flag-Rab40b, or Flag-Rab40b-4A cells on collagen coating, plated in either low or high serum conditions. Results are of three separate runs, at least 3 wells per condition per run. Left panel; Grouped scatter plot of relative migration over entire time-course. Right panel; Bar graph showing relative migration at 24 hours. (F) Chemotactic assays of control or Flag-Rab40b-4A cells plated on either collagen or fibronectin. Results are of three separate runs, at least 3 wells per condition per run.

General observations from our live cell imaging suggested differences in the actin cytoskeleton structure and dynamics, which are known to play a vital role in cell migration and invasion. Since Cullin family members, including Cullin5, have been shown to exert effects on the actin cytoskeleton [41, 47–49], we next sought to assess changes to the cytoskeleton that result from diminished Rab40b/Cullin5 binding. As shown in Figure 2C-D, expression of Flag-Rab40b-4A enhanced formation of ventral actin stress fibers compared to control cells or cells expressing wild-type Flag-Rab40b. Stress fibers are contractile actomyosin bundles that contribute to mechanical force generation to regulate cell contractility, adhesion and motility [50]. Thus, the inability of cells to properly reorient or maintain their actin cytoskeletal architecture could explain why Flag-Rab40b-4A expressing cells can still move but have difficulties sustaining directional movement.

### Rab40b/Cullin5 Regulates Chemotactic Migration and Cell Invasion

Having observed defects in cell directionality and cytoskeletal makeup in Flag-Rab40b-4A cells, we hypothesized that Rab40b and Cullin5 binding may govern more complex modes of cellular movement. A defining feature of SOCS family member proteins is the ability to regulate cellular responses to extracellular signaling cues, so we next assessed how the loss of Rab40b/Cullin5 binding impacted chemotactic migration. Serum-starved control, wild-type Flag-Rab40b and Flag-Rab40b-4A cells were seeded in low-serum containing chambers and allowed to migrate to chambers containing complete medium as a chemo-attractant. As shown in Figure 2E, Flag-Rab40b-4A cells demonstrated a significant reduction in migration toward chemoattractant. As a control, we also plated cells in chambers containing high-serum. In these conditions both control and wildtype Flag-Rab40b cells, in addition to Flag-Rab40b-4A cells, exhibit a reduced ability to migrate toward serum. These results suggest that movement by control and Flag-Rab40b cells in the low-serum condition is driven by chemotaxis and that Flag-Rab40b-4A expressing cells are deficient in this ability.

Extracellular substrates can alter integrin signaling and differentially effect cell migration on particular substrates [51], so we next asked if different surface substrates would affect chemotactic migration of Flag-Rab40b-4A cells. Figure 2F demonstrates that reduced chemotactic migration of Flag-Rab40b-4A cells occurs on either collagen or fibronectin, suggesting that the Rab40b/Cullin5 complex may be part of a core chemotactic migration machinery rather than mediating a response to specific extracellular matrix components. Cell proliferation was not affected in our mutant cell line (Supplemental Figure 1E), suggesting that the differences we observe in chemotactic migration are not due to inherent differences in cell division.

Next, we sought to analyze how Rab40b and Cullin5 binding impacts cell migration in a three dimensional (3D) ECM environment. Using a modified inverted 3D Matrigel invasion assay, we found that while expressing wild-type FLAG-Rab40b had no significant impact on cell invasion compared to control cells, as expected [18], cells expressing the FLAG-Rab40b-4A mutant had a significantly reduced ability to migrate through a 3D Matrigel matrix (Figure 3A-B). Degradation and remodeling of the ECM during cell migration is facilitated by protrusive actin structures called invadopodia. Our lab has previously demonstrated that Rab40b is required for the formation and function of invadopodia by regulating the secretion of MMP2 and MMP9. Given our current results showing a Rab40b/Cullin5 binding mutant induces changes to the actin cytoskeleton and impairs the cells ability to migrate through an extracellular environment, we hypothesized that the Rab40b/Cullin5 complex may function by co-regulating MMP secretion with actin remodeling during invadopodia formation and extension. To test this hypothesis, we stained our cell lines with phalloidin-Alexa594 along with an anti-cortactin antibody, an established marker of maturing invadopodia [52, 53], and then counted the number of cortactin-positive actin puncta per cell. Consistent with our previously published data [17], overexpression of wild-type Rab40b had no effect on invadopodia number (Figure 3C-D). However, cells expressing Flag-Rab40b-4A show a significant decrease in the number of invadopodia per cell (Figure 3CD), suggesting that the ability of Rab40b to promote invadopodia formation depends on CRL5 binding. Since cortactin is also known to be present on late endosomes and lysosomes we co-stained cells with cortactin and CD63, a well-established marker for endo-lysosomal pathway. As shown in Supplemental Figure 3A, expression of Flag-Rab40b-4A had no effect on the number of cortactin and CD63 positive organelles, suggesting that Rab40b-4A mutant affects invadopodia formation rather than lysosomal. Given that our cells show a decrease capacity to migrate through a 3D EMC environment, coupled with less invadopodia, we surmised that Flag-Rab40b-4A cells may be secreting less matrix degrading enzymes. Previous work from our lab has demonstrated the necessity of Rab40b in regulating secretion of MMP at invadopodia [17], so we next assessed the level of secreted MMP2 and MMP9 from our cells. In accordance with less invadopodia, the Flag-Rab40b-4A cell line shows decreased secretion of MMP9 in a 2D gel zymography assay as compared to control and wild-type Flag-Rab40b expressing MDA-MB-231 cells (Figure 3E). This suggests that the ability of Rab40b to regulate MMP secretion is dependent on Rab40b binding to Cullin5, and that our observed decrease in cell invasion through an extracellular matrix can be explained in part by the decreased secretion of matrixdegrading enzymes.

**Figure 3.**
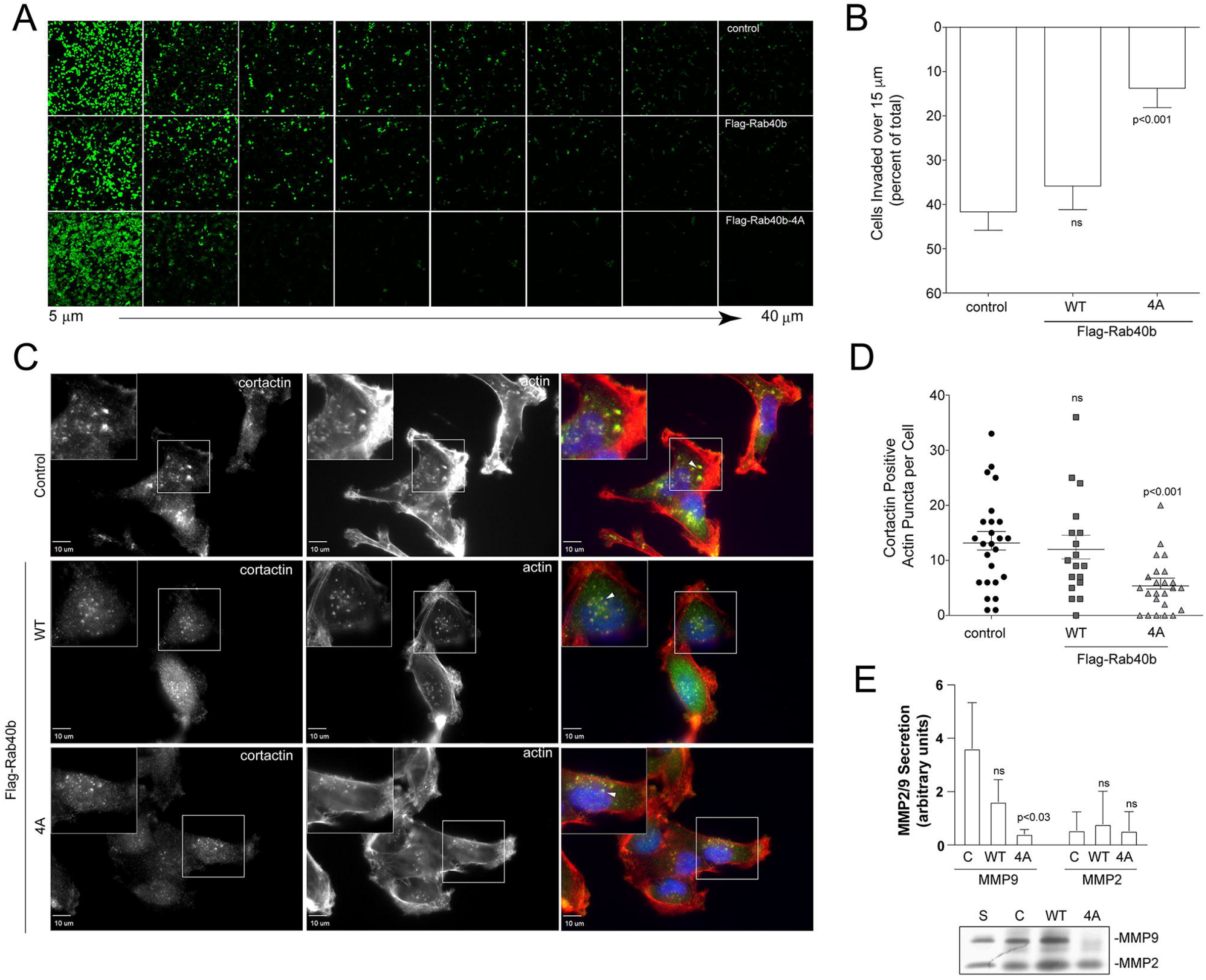
Rab40b/Cullin5 Regulates Chemotactic Migration and Cell Invasion. (A-B) Matrigel invasion assay of control, Flag-Rab40b and Flag-Rab40b-4A cells. Representative field-of-view images of Calcein-stained cells at 5 μm increments throughout the plug. (B) Quantification of cell migration through Matrigel plug. Results are of three separate runs, at least three fields-of-view per condition per run. (C-D) Control, Flag-Rab40b or Flag-Rab40b-4A cells plated on collagen coated coverslips, fixed and stained with phalloidin-594 (red), cortactin (green) and DAPI (blue). Inset regions of interest highlight phalloidin/cortactin dual-positive puncta. Arrows point to actin/cortactin puncta. (D) Quantification of dual-positive actin/cortactin puncta. n ≥ 18 cells per condition. (G) Gelatin zymography analysis of MMP2 and MMP9 secretion from control, Flag-Rab40b and Flag-Rab40b-4A cells grown in serum-free media for 24 hours. Fetal bovine serum (serum) contains MMP2/9 and was used as a positive control. The data shown are the means and standard error of means derived from three independent experiments. The data were normalized against serum (S) levels of MMP2 and MMP9.

### Rab40b/Cullin5 Regulates Localization and Dynamics of Focal Adhesion Sites

Stress fibers are known to connect to FAs and regulate FA dynamics [54]. Furthermore, FA dynamics has been shown to be regulated by Cullin5 [44]. In order to assess whether the Flag-Rab40b-4A-dependent increase in stress fibers was accompanied by an increase in FAs we next stained MDA-MB-231 cells with antibodies against the FA marker paxillin. As shown in Figure 4A, C-D, we see an increase in both number and size of FAs in the Flag-Rab40b-4A mutant cells. Western Blot analysis also shows an increase in paxillin protein levels in Flag-Rab40b-4A cells compared to control and Flag-Rab40b lines (Figure 4B). To determine whether the increase in FA number was due to an increase in their stability, we transfected either control or Flag-Rab40b-4A cells with GFP-paxillin and SiR-actin and imaged them every 4 minutes for up to 7 hours (Supplemental Movie 3 and Supplemental Movie 4). Images were then uploaded to the FAAS server to assess FA lifespan. As shown in Figure 4E, focal adhesions are longer lived in Rab40b-4A mutant cells. Additionally, Rab40b-4A mutants have a greater percentage of adhesions that are further from the cell boundary (Figure 4F), a result consistent with more stable, less dynamic adhesions [55]. Consistent with this, FAs in Flag-Rab40b-4A cells were more likely to be zyxin positive (Supplemental Figure 2B-C), a marker for more mature focal adhesions [56], again suggesting FAs in Flsg-Rab40b-4A-expressing cells are more stable than in their control counterparts.

**Figure 4.**
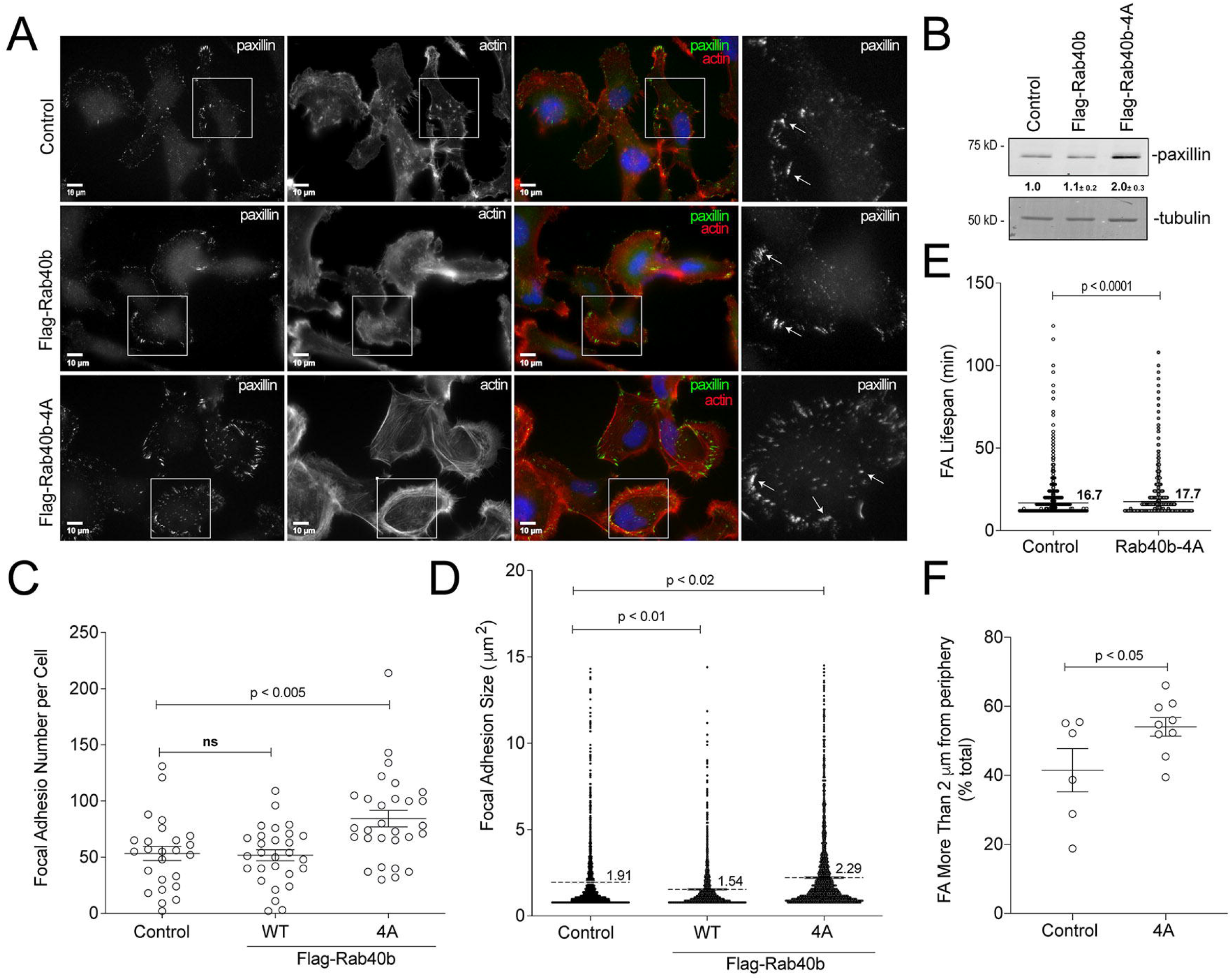
Rab40b/Cullin5 regulates localization and dynamics of Focal Adhesion sites. (A) Representative images of control, Flag-Rab40b, and Flag-Rab40b-4A MDA-MB-231 cells fixed and stained with phalloidin-Alexa594 (red), anti-paxillin (green, focal adhesion marker) and DAPI (blue). Insets highlight anti-paxillin stained regions of interest. (B) Western Blot analysis of cell lysates using anti-paxillin (top blot) and anti-tubulin (bottom blot) antibodies. Numbers shown are densitometry analysis from at least 3 separate experiments, relative to tubulin and standardized to control levels. Control::Flag-Rab40b-4A p < 0.05. (C) Quantification of number of focal adhesions per cell for control, Flag-Rab40b and Flag-Rab40b-4A cells. n ≥ 20 cells per condition. (D) Quantification of focal adhesion size in control, Flag-Rab40b, and Flag-Rab40b-4A cells. n=2479 total adhesions analyzed. (E) Quantification of focal adhesion average lifespan in control and Flag-Rab40b-4A cells. Average lifespan of focal adhesions was calculated from time-lapse images, 3 cells per condition. (F) Quantification of percent of focal adhesions within 2 μm of cell border in control and Flag-Rab40b-4A cells.

Focal adhesion kinase (FAK) regulates FA formation and dynamics. Thus, we hypothesized that differences in FAK signaling may contribute to differences in FA dynamics in Flag-Rab40b-4A cells. We assessed the level of two different FAK phosphorylation sites: Y397 and S910. Y397 is an autophosphorylation site that enables binding of Src, FAK activation, and promotes focal adhesion stability [57, 58]. In contrast, S910 has been implicated in stimulating paxillin binding and turnover [59], and is necessary for invasive migration [60]. As shown in Supplemental Figure 2D, in the Flag-Rab40b-4A cell line, we see an increase in pFAK Y397 and a decrease in pFAK S910 compared to control and wild-type expressing cells, while total FAK remains constant. This result is consistent with the hypothesis that our observed increase in focal adhesions stability is due, in part, to differences in FAK signaling.

So far, our data demonstrate that expression of Flag-Rab40b-4A mutant leads to changes in size, number and subcellular distribution of FA. To test whether blocking Rab40b and Cullin5 binding also leads to changes in cell-ECM adhesion we plated control and Flag-Rab40b-4A cells on collagen coverslips and allowed them to adhere and spread for 90 minutes. Cells were then fixed and surface area of spreading visualized and analyzed by staining with phalloiding-Alexa594. As shown in Supplemental Figure 5A, Flag-Rab40b-4A cells spread faster as compared to control cells, indicating that increase in focal adhesions function to adhere the cells to the substrate.

### EPLIN is a Rab40b Binding Proteins that is Ubiquitylated by Rab40/Cullin5 Complex

We next set out to identify the substrates that may be targeted by this complex. Since SOCS box containing scaffolds recruit proteins for ubiquitylation, we first tested whether known Rab40b bound proteins, such as Tks5 [18], are degraded in a Rab40b/Cullin5 dependent fashion. To determine that we analyzed whether overall cellular levels of Tks5 are increased in cells over-expressing Flag-Rab40b-4A mutants, since that would be expected if Rab40b/Cullin-5 mediates Tks5 ubiquitylation and degradation. Surprisingly, the Flag-Rab40b-4A mutant had little effect on total Tks5 levels compared to control cells (Supplemental Figure 3B), suggesting that Tks5 may not be a substrate for Rab40b/Cullin5 complex ubiquitylation. We next tested if p130Cas, a known FA protein [61–63] that has been shown to be regulated by Cullin5 [48], was affected by loss of Rab40b/Cullin5 binding. As shown in Supplemental Figure 3B, p130Cas is also not stabilized with loss of Rab40b/Cullin5 complex formation, suggesting that the Rab40b/Cullin5 complex may act on as of yet unknown set of proteins.

Typically, upon ubiquitylation substrate proteins will rapidly dissociate from the E3 ligase complex, making identification of specific target proteins very difficult. We surmised that certain Rab40b-dependent substrates will remain bound to Rab40b if Cullin5 binding is blocked or diminished, thus allowing us to identify them by mass spectrometry. To that end, we harvested lysates from cells expressing either wild-type Flag-Rab40b or mutant Flag-Rab40b-4A, immunoprecipitated Flag-Rab40b with anti-Flag antibodies and identified bound proteins by mass spectrometry. As shown in Figure 5A, Cullin5, Elongin B and Elongin C coimmunoprecipitation with Flag-Rab40b-4A was diminished compared to wild-type Flag-Rab40b. We also pulled out Rbx2 (known also as Rsf7 or Sag1), a known component of CLR complexes (Figure 5A), as a Rab40b/Cullin5 binding protein. To define putative Rab40b/Cullin5 substrates we next filtered all candidates to focus on proteins that were enriched more than 2-fold in the Flag-Rab40b-4A sample compared to wild type Flag-Rab40b (Figure 5A).

**Figure 5.**
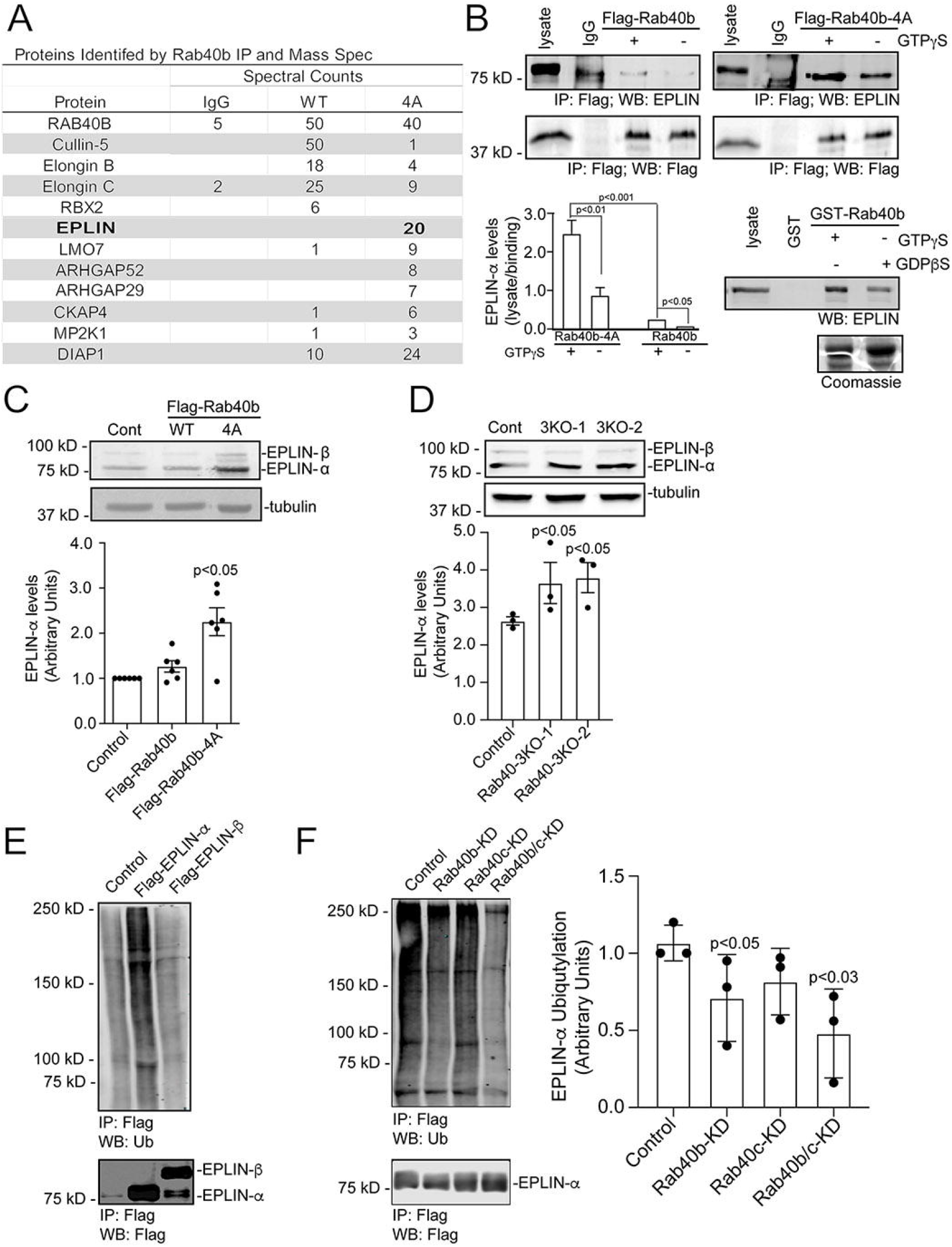
Rab40b/Cullin5 regulates EPLIN stability and localization. (A) Abbreviated list of proteins identified by immunoprecipitation of Flag-Rab40b and Flag-Rab40b-4A cells, followed by mass spectroscopy analysis. (B) Rab40b and EPLIN interaction analysis. Top panels; Cell lysates of Flag-Rab40b and Flag-Rab40b-4A cells in the presence or absence of GTPγS were incubated with either IgG or an anti-Flag antibody and immunopreciptated with protein G beads. Immunoprecipitates were analyzed by Western Blot with anti-Flag and anti-EPLIN antibodies. Bottom left; Quantification of EPLIN and Flag-Rab40b binding. Data are the means and SEM derived from three independent experiments. In all cases signal was normalized to the EPLIN signal in lysate. Bottom right; MDA-MB-231 control cell lysates were incubated with glutathione beads coated with either GST or GST-Rab40b, in the presence of GDPβS or GTPγS. Bound EPLIN was eluted and analyzed by Western Blot. (C) Western Blots of endogenous EPLIN in control, Flag-Rab40b, and Flag-Rab40b-4A cells. Quantification below are the means and SEM derived from three different experiments and normalized against tubulin levels. (D) Western Blots of endogenous EPLIN in control and Rab40a/b/c-KO (3KO) cells. Two different 3KO lines were used in these experiments. Quantification below are the means and SEM derived from three different experiments and normalized against tubulin levels. (E) Western Blot images of 293t cells transfected with empty vector control (CNT), Flag-EPLIN-α or Flag-EPLIN-β for 24 hours, treated with 10 μm MG132 overnight, harvested and immunoprecipitated for Flag, then blotted for either ubiquitin (top) or Flag (bottom). (F) Western Blot images of 293t cells treated with siRNAs for non-targeting control (siCNT), Rab40b, Rab40c or both Rab40b and Rab40c, transfected with Flag-EPLIN-α, followed by treatment with MG132 overnight, harvested and immunoprecipitated for Flag, then blotted for either ubiquitin (top) or Flag (bottom). Quantification on the right are the means and SEM derived from three different experiments.

One highly enriched candidate was EPLIN (*E*pithelial *P*rotein *L*ost *I*n *N*eoplasia; also referred to as LIMA1), an actin bundling protein that is known to be a negative regulator of cell migration [36]. To test if binding of EPLIN to Rab40b occurs in a GTP-dependent fashion, we again immunoprecipitated Flag from cell lysates expressing wild-type or mutant Flag-Rab40b in the presence or absence of non-hydrolysable GTP analog GTPγS, and then probed for binding to EPLIN by Western Blot. As shown in Figure 5B, EPLIN binds poorly to wild-type Flag-Rab40b, consistent with our proteomic analysis. Importantly, Rab40b-4A mutation substantially increased EPLIN’s ability to bind to Rab40b. Finally, Rab40b and EPLIN binding is enhanced by GTPγS (Figure 5B). To further confirm that EPLIN binds to Rab40b in GTP-dependent fashion we next incubated control MDA-MB-231 lysates with glutathione beads coated with purified GST-Rab40b pre-loaded with either GTPγS or GDPβS. Western Blot analysis of eluted proteins again shows that EPLIN binds with greater affinity to GTP-Rab40b (Figure 5B). This data demonstrates that the nucleotide state of Rab40b regulates the binding of EPLIN, and thus supports the conclusion that EPLIN is a canonical Rab40b effector.

While our data show that EPLIN binds to Rab40b in a manner consistent with classical Rab effectors [64], its enhanced binding to a Rab40b-4A mutant also suggests that EPLIN might be subject to Rab40b/CRL5 dependent ubiquitylation, and that this ubiquitylation may diminish EPLIN ability to interact with Rab40b. To test this hypothesis, we first transfected 293t cells with either Flag-tagged EPLIN-α or EPLIN-β, immunoprecipitated with anti-Flag antibodies, and Western Blotted for ubiquitin. As shown in the Figure 5E, EPLIN-α was ubiquitylated, suggesting that Rab40b/Cullin5 may specifically regulate ubiquitylation of EPLIN-α in these conditions. While we did not detect ubiquitylation of Flag-EPLIN-β we cannot discount the possibility that the sensitivity of our assays was not good enough to detect EPLIN-β ubiquitylation or that 293t cells may not recapitulate the signaling environment needed to ubiquitylate EPLIN-β, especially since 293t cells are not migratory cell type. To test if Rab40b is targeting Flag-EPLIN-α for ubiquitylation, 293t cells were transfected with Flag-EPLIN-α and siRNA for Rab40b. Lysates were then immunoprecipitated with anti-Flag antibodies and blotted for ubiquitin. Surprisingly, depletion of Rab40b (Supplemental Figure 3C) caused only slight decrease in ubiquitylated EPLIN-α (Figure 5F). It has been shown that closely related Rab family members can bind the same effector proteins and have overlapping functions [64–67]. Since most mammalian cells express Rab40c, including MDA-MB-231 cells, we also used siRNA to knock-down Rab40c alone as well as both Rab40b and Rab40c in 293t cells (Supplemental Figure 3C) and similarly assessed EPLIN-α ubiquitylation. While loss of Rab40c had no effect, loss of both Rab40b and Rab40c isoforms together caused a further decrease in the ubiquitylation of EPLIN-α (Figure 5F). This supports the conclusion that Rab40b can mediate EPLIN ubiquitylation and that other Rab40 isoforms, such as Rab40c, may also contribute to this process.

If Rab40b-dependent EPLIN ubiquitylation targets EPLIN for degradation, then it would be expected that expression of Flag-Rab40b-4A should lead to stabilization of EPLIN in cells. Consistent with this hypothesis, Western Blot analysis shows total levels of EPLIN are significantly increased in the Rab40b-4A mutants (Figure 5C) compared to control and wild type Flag-Rab40b-expressing cells. To further test the hypothesis that Rab40 may mediate EPLIN ubiquitylation and degradation we next generated CRISPR-mediated knock-out of three Rab40 family members Rab40a, Rab40b and Rab40c (Rab40-3KO). As shown in Figure 5D, knock-out of all three Rab40 isoforms show a similar increase in EPLIN-α compared to control cells. This suggests that in addition to Rab40b, other members of the Rab40 family may compensate for loss of Rab40b in ubiquitylating EPLIN. Thus, in rest of the study we used Ran40-3KO for all functional studies.

### Rab40/Cullin5 Affects Actin Cytoskeleton by Regulating Sub-Cellular EPLIN Distribution

EPLIN targeting to specific subcellular domains was shown to play a key role in regulating EPLIN function [68]. As shown in Figures 6A, immunofluorescent staining with a pan-EPLIN antibody shows that in control MDA-MB-231 cells EPLIN is localized at the lamellipodia, just behind lamellipodia actin ruffles (also see figure 7E). This is consistent with the proposed function of EPLIN: regulation of actin bundling during formation of actomyosin stress fibers that are localized just behind leading edge of lamellipodia and are required for directional cell migration (Figure 7A) [69]. In contrast, in Flag-Rab40b-4A and Rab40-3KO cells EPLIN strongly associates with stress fibers (Figure 6B-C), and was present at the leading edge of the lamellipodia (Figure 7B-G). Interestingly, lamellipodia in Flag-Rab40b-4A and Rab40-3KO cells were smaller and had less pronounced actin ruffles, an observation consistent with reports that EPLIN-induced actin filament bundling may inhibit Arp2/3-dependent actin ruffling [68, 70].

**Figure 6.**
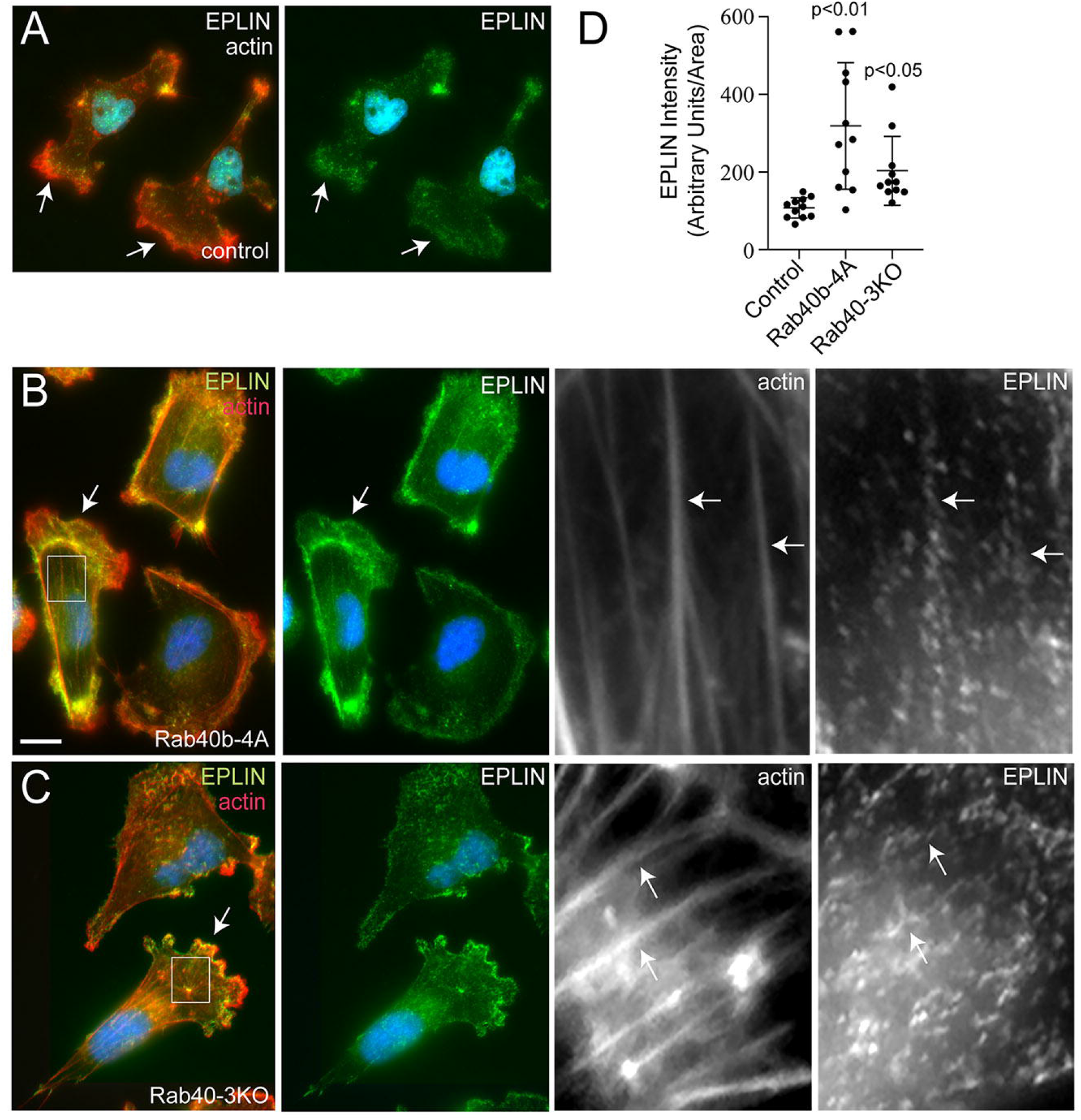
Rab40a/b/c depletion or Rab40b-4A overexpression leads to increase in plasma membrane and stress-fiber associated EPLIN. (A-C) Immunofluorescent images of control (A), Flag-Rab40b-4A (B) and Rab40-3KO (C) cells plated on collagen coated coverslips, fixed and stained with anti-EPLIN antibodies (green) and phalloidin-Alexa594 (red). Boxes mark regions of interest. Arrows in whole field images indicate lamellipodia leading edge. Arrows in boxed regions indicate stress fibers. (D) Quantification of EPLIN fluorescence in control, Flag-Rab40b-4A and Rab40-3KO cells. The data shown are the means and SEM derived from three different experiments. Dots represent individual cells analyzed.

**Figure 7.**
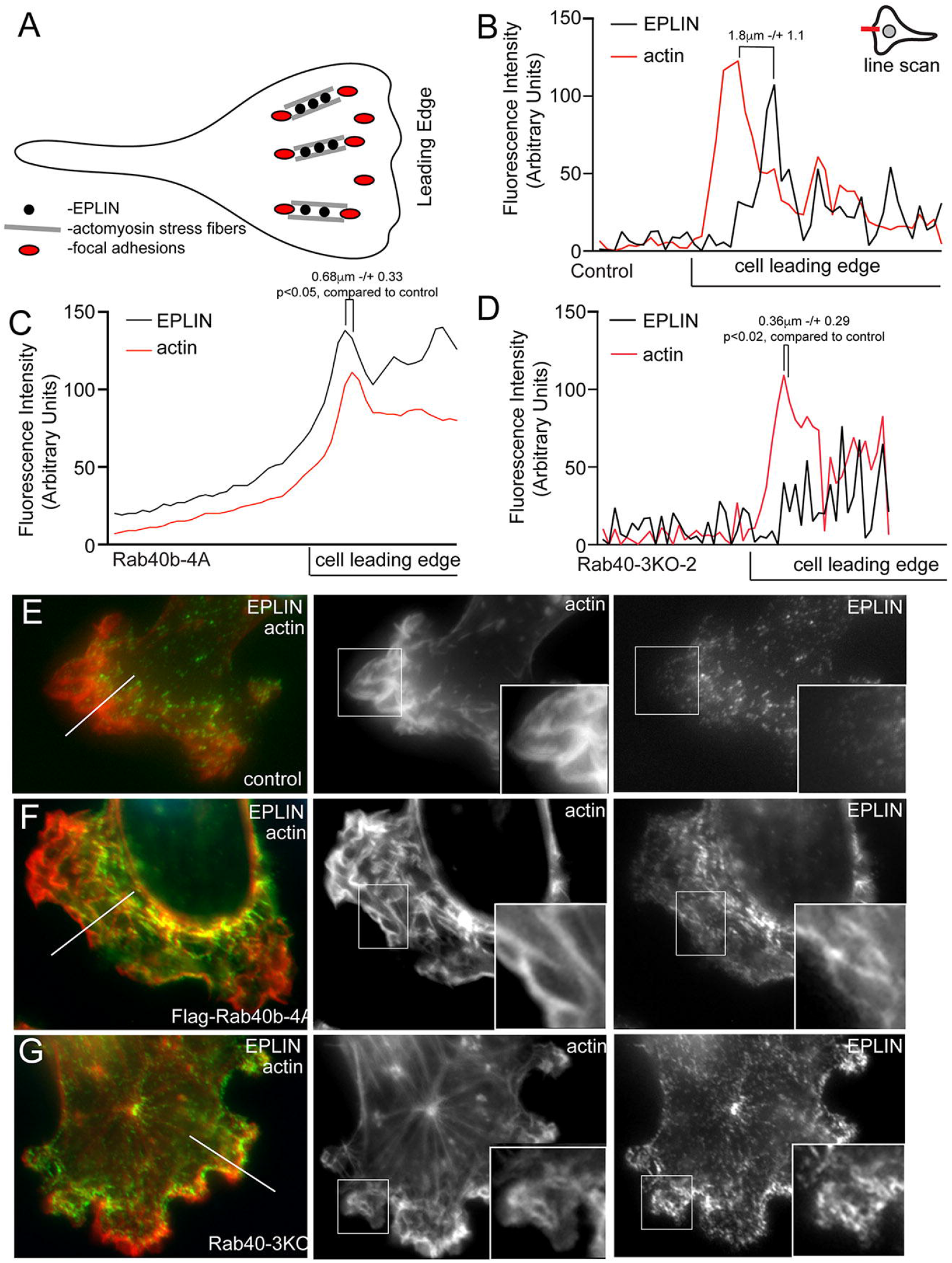
Rab40b inhibits EPLIN accumulation at the lamellipodia leading edge. (A) Schematic representation of FA and stress fiber distribution in lamellipodia. (B-G) Representative line scans from control (E), Flag-Rab40b-4A (F), and Rab40-3KO (G) cells plated on collagen-coated coverslips, fixed and stained with phalloidin-Alexa594 (red) and anti-EPLIN (green) antibodies. Boxes mark the region of interest shown in inset. Line marks the region analyzed by line-scan (quantifications shown in B-D). Eplin distance from actin front; control, 1.8 μm ± 1.1; Flag-Rab40b-4A, 0.68 μm ± 0.33 (p<0.05); and Rab40-3KO 0.36 μm ± 0.29 (p<0.02). n = 5 cells for each cell line.

Because recent work has suggested differences in localization and function for each EPLIN isoform [68, 71] we next asked if the subcellular localization changes of EPLIN we observed were due to differential regulation of individual isoforms. We transfected control or Flag-Rab40b-4A cells with GFP-EPLIN-α or GFP-EPLIN-β. Although stress fibers are generally absent in control MDA-MB-231 cells, both isoforms are present along the stress fibers induced by Flag-Rab40b-4A mutant cells [68]. Additionally, in control cells GFP-EPLIN-β localizes just behind the actin rich front edge of lamellipodia (Supplemental Figure 6A). Similar to endogenous GFP-EPLIN distribution, in Rab40b-4A cells GFP-EPLIN-β no longer lags behind the actin ruffles in lamellipodia, which suggest that Rab40b/Cullin5 binding influences localization of EPLIN-β during cell migration.

### Rab40/Cullin5 Regulates Lamellipodia Dynamics during Cell Migration

We next sought to analyze the subcellular distribution of Rab40b to better understand where the Rab40b/Cullin5 complex may function to regulate EPLIN ubiquitylation and degradation. To that end, we generated MDA-MB-231 cell lines stably expressing either GFP-Rab40b or GFP-Rab40b-4A. As shown in Figure 8A, while the majority of GFP-Rab40b is present in the cytosol, a sub-population of GFP-Rab40b can clearly be observed at the lamellipodia where it colocalizes with actin ruffles. GFP-Rab40b-4A is also present at the front end of lamellipodia and colocalizes with actin, suggesting that inhibition of Cullin5 binding does not affect sub-cellular localization of Rab40b (Figure 8B). As was the case with our aforementioned data, GFP-Rab40b-4A expressing cells had diminished levels of actin ruffles (Figure 8B) and an increase in FAs compared to the cells expressing wild-type GFP-Rab40b (Figure 8C-D). Similarly, in GFP-Rab40b-4A cells EPLIN accumulates at the leading edge of lamellipodia where it colocalizes with GFP-Rab40b-4A, an association largely absent in cells expressing wild-type GFP-Rab40b (Figure 8E-F).

**Figure 8.**
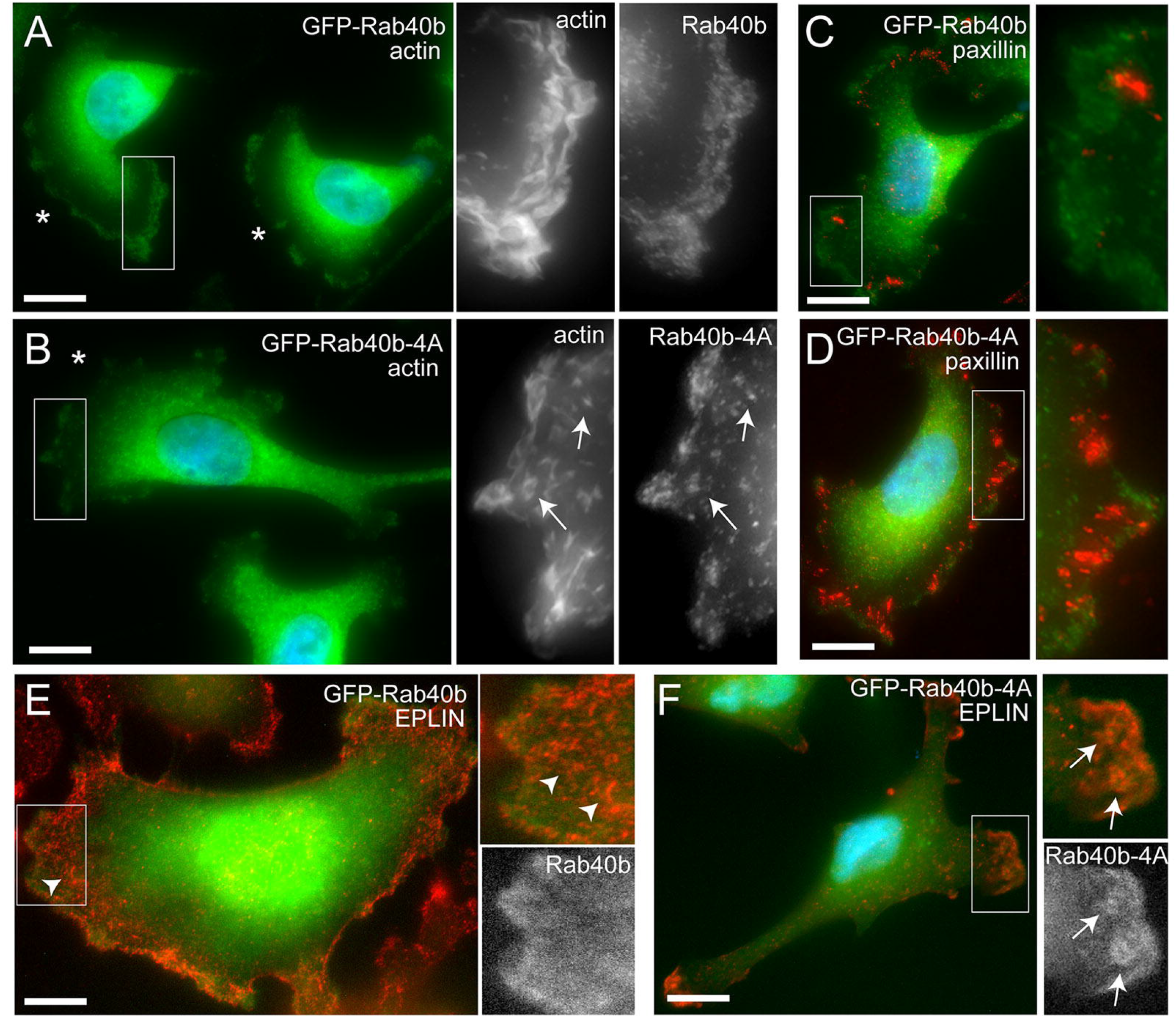
GFP-Rab40b co-localizes at the actin ruffles at the leading edge of lamellipodia. (A-F) Representative images of GFP-Rab40b (A, C and E) or GFP-Rab40b-4A (B, D, and F) expressing MDA-MB-231 cells plated on collagen-coated coverslips, fixed and stained with phalloidin-Alexa594 (red, A-B), anti-paxillin (C-D) or anti-EPLIN antibodies (E-F). Boxes mark the region of interest shown in inset. Asterisks mark leading edge of lamellipodia. Arrowheads mark EPLIN staining that does not colocalize with GFP-Rab40b. Arrows point to structures positive for both, EPLIN and GFP-Rab40b-4A.

EPLIN has a well-established role in inhibiting Arp2/3 branched actin polymerization [68, 70], a process known to be essential for actin ruffling at the leading edge of cells. The accumulation of EPLIN and altered actin ruffles in our Rab40b mutant cells suggests altered lamellipodia dynamics. To that end we imaged lamellipodia plasma membrane dynamics using DIC time-lapse microscopy. As expected, control cells exhibited very active lamellipodia ruffling (Supplemental Figure 5C, Supplemental Movie 5). In contrast, overexpression of FLAG-Rab40b-4A or depletion of Rab40a/b/c (Rab40-3KO) led to decrease in lamellipodia ruffling (Supplemental Figure 5C, Supplemental Movies 6-7). Together, this data suggests that Rab40b mediates removal of EPLIN at the leading edge of migrating cells, whereas expression of mutant Rab40b-4A inhibits EPLIN ubiquitylation, thus, stabilizing Rab40b and EPLIN interaction and leading to the accumulation of Rab40b/EPLIN complexes at the leading edge of lamellipodia and stress fibers (see proposed model in Figure 10). Importantly, Rab40-3KO phenocopies the effects of over-expressing Flag-Rab40b-4A, where we observe an increased number of FAs (Supplemental Figure 4E), stimulation of stress fibers (Supplemental Figure 3D) and increased cell-ECM adhesion (Supplemental Figure 5B). If depletion of Rab40a/b/c or overexpression of Rab40b-4A stabilizes stress fibers, one would predict that that should also lead to increase in stress fiber-associated non-muscle Myosin IIA/B. We therefore stained MDA-MB-231 cells with anti-nonmuscle Myosin IIA/B antibodies. Consistent with our hypothesis, Flag-Rab40b-4A and Rab40-3KO cells exhibited increase in stress fiber associated non-muscle Myosin IIA/B (Supplemental Figure 4A-C).

Overall, our results suggest that Rab40b complexes with Cullin5 to regulate sub-cellular EPLIN localization by decreasing EPLIN levels at the leading edge of lamellipodia and allowing EPLIN accumulation at stress fibers and invadopodia. It is likely, however, that EPLIN is only one of several Rab40b substrates and that Rab40b/Cullin5 regulates the activity of multiple actin regulators. Accordingly, our proteomic screen identified several other regulators of actin dynamics that are putative substrates for the Rab40b/Cullin5 complex (Figure 5A, see Discussion).

### Rab40b/Cullin5 Regulates Primary Tumor Growth *In Vivo*

To investigate the function of Rab40b/Cullin5 binding *in vivo*, we performed mammary fat pad injections into athymic nude mice with control, wild-type Flag-Rab40b or mutant Flag-Rab40b-4A expressing MDA-MB-231 cell lines. At 60 days post-injection, we observed severely reduced tumor growth in the Rab40b-4A injected animals (Figure 9A) and all Rab40b-4A injected mice survived to the end of the 8-week study. Conversely, tumor burden necessitated the mice with control and wild-type tumors be sacrificed at earlier time-points (Figure 9B).

**Figure 9.**
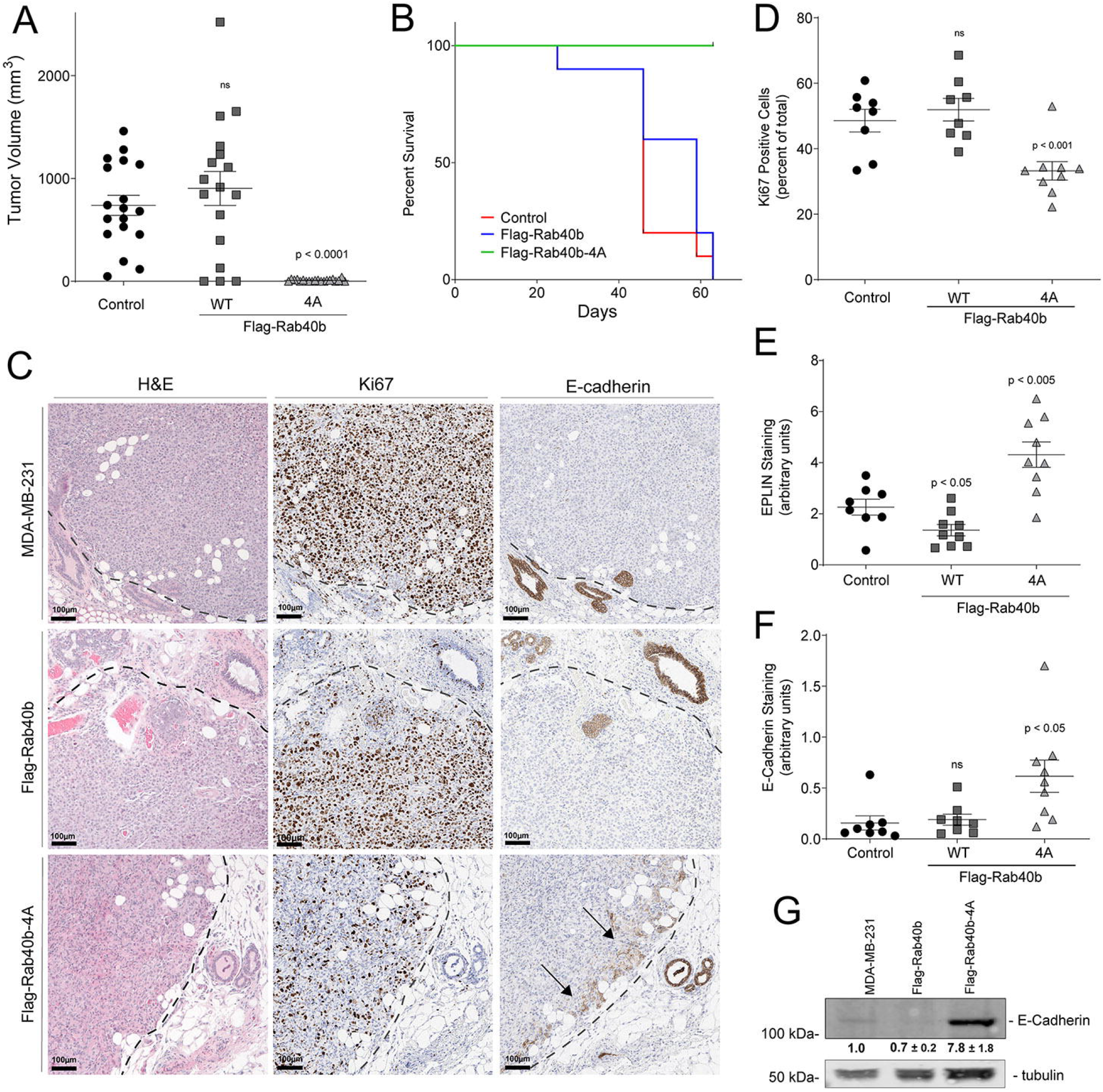
Rab40b/Cullin5 co-regulates primary tumor growth in vivo. (A) Tumor volumes from xenograft assays. Control - 739 cm^3^ ± 414, n=18; Flag-Rab40b - 904 cm^3^ ± 682, n=17; Flag-Rab40b-4A - 8.9 cm^3^ ± 10, n=20 tumors. (B) Kaplan-Meier survival plot showing overall survival of mice groups throughout study. n=10 mice per group. (C) Representative images of immunohistological stains of serial sections from tumors grown from each cell line. Dashed lines indicate tumor border. Arrows highlight increased E-cadherin staining at tumor periphery. (D) Quantification of Ki67 staining as a percentage of positive pixels from tumors represented in (C). Control - 48.6 ± 9.8, n = 8; Flag-Rab40b - 51.9 ± 9.8, n = 8; Flag-Rab40b-4A - 33.2 ± 8.4, n = 9. (E) Quantification of strong EPLIN stains as a percent of positive pixels from tumors represented in (C). Control - 2.3 ± 0.9 SEM, n = 8; Flag-Rab40b - 1.4 ± 0.7 SEM, n = 8; Flag-Rab40b-4A - 4.4 ±1.5 SEM, n = 9. (F) Quantification of E-Cadherin stains as a percentage of positive pixels from tumors imaged in (C). Control - 15.6 ± 6.9 SEM, n= 8; Flag-Rab40b - 18.9 ± 5.3 SEM, n=8; Flag-Rab40b-4A - 61.6 SEM ± 15.9 SEM, n = 9. (G) Western Blot of E-Cadherin from cell lysates of control, Flag-Rab40b and Flag-Rab40b-4A cells. Control::4A p < 0.05. Numbers shown are the average densitometry analysis derived from at least three biological replicates, relative to tubulin and normalized to control levels.

Immunohistochemical analysis of a proliferation marker, Ki67, shows a significant decrease in staining throughout Flag-Rab40b-4A mutant tumors compared to both control and wild-type Flag-Rab40b expressing cells (Figures 9C-D), suggesting that the loss of Rab40b and Cullin5 decreases cell proliferation to result in decreased tumor growth. We next asked if increased EPLIN we observe *in vitro* would persist in *in vivo*. We observed a significant increase in EPLIN staining in the Flag-Rab40b-4A tumors (Figure 9C and E; Supplemental Figure 6E).

E-cadherin is another well-known intercellular adhesion molecule that inhibits the migratory capability of cells by enhancing cell-cell adhesion and inhibiting epithelial-to-mesenchymal transition. Furthermore, Cullin-5 has been implicated in E-cadherin degradation [72]. We observe that primary tumors derived from Flag-Rab40b-4A mutant cells show increased E-cadherin expression, with a particular subset of cells having stronger staining along the tumor border (Figure 9C, F). Furthermore, increased E-cadherin expression is mirrored in the MDA-MB-231 cell lines engineered to overexpress the Flag-Rab40b-4A mutant (Figure 9G).

## DISCUSSION

The coupled acts of cell migration and invasion involve very complex and highly interconnected molecular pathways that must be properly coordinated to ensure correct organism development and function. Previous reports from our lab have identified Rab40b as an important protein for the secretion of MMPs at invadopodia and central for cell invasion and migration [7, 17, 18]. Rab40b belongs to a unique Rab40 sub-family of small monomeric GTPases that contain SOCS domain at their C-terminus, just before geranylgeranylation site. SOCS domains are known to mediate interaction with CRLs, thus, we hypothesized that Rab40b may mediate its function by regulating protein ubiquitylation. While it has been shown that Rab40a and Rab40c bind Cullin5, it remains unclear whether the same is true for Rab40b. Here we sought to further understand how Rab40b impacts other aspects of cell invasion in addition to MMP secretion by exploring two open questions: does Rab40b/Cullin5 binding play a role in regulating cell migration and invasion, and what are the substrates of this complex?

We show that in breast cancer cells Rab40b does bind to the CRL5 E3 ligase complex containing Cullin5, Elongin B, Elongin C, and Rbx2 and that mutations to the Rab40b-SOCS box are sufficient to disrupt this interaction. We also demonstrate that loss of the Rab40b/Cullin-5 complex by Rab40b-4A mutation impacts chemotactic migration and invasion. Analysis of individual cell movement also revealed defects and alterations in cell behavior. While cell speed remains unchanged, there are noticeable differences in cell directionality, which suggests that the deficiency of Rab40b-4A-expressing cells to move toward a chemoattractant may be due to a loss in directionality. Mutant cells also have less invadopodia and, accordingly, secrete less of the ECM degrading protein MMP9. This would explain, in part, the inability to move through a 3D ECM environment. Interestingly, our time-lapse analyses also suggest that Rab40b-4A cells have increased cell-cell and cell-ECM adherence, thus, leading to formation of cell clumps and apparent loss of Contact Inhibition of Locomotion. In agreement with this hypothesis, we observe that Flag-Rab40b-4A cells have a noticeable increase in the cell junction protein E-cadherin, a purported Cullin5 target [72], compared to control and wild-type Flag-Rab40b expressing cells, which suggests that mutant cells may remain connected and could be thus less efficient at migrating through 3D Matrigel.

Mutant Rab40b-4A and Rab40-3KO cells also adhere stronger to the ECM and exhibit an increase in ventral stress fibers. Previous reports have shown that stress fibers can be induced in MDA-MB-231 cells by activating RhoA [73, 74], suggesting that Rab40b-4A cells may have altered RhoA signaling. We also observe an increase in both the number and size of focal adhesions (FAs) per cell, with an accompanying increase in total paxillin, as well as more zyxin positive FAs. This suggests that FAs are more stable in mutant Rab40b-4A and Rab40-3KO cells. Indeed, live-imaging analysis shows that adhesions have a greater life-span in Rab40b-4A cells. It was recently shown that Rab18 regulates focal adhesion dynamics in U2OS cells. Rab18 is closely related to the Rab40 family [75], and it has been suggested that Rab40 split from the Rab18 family and gained the SOCS-box upon the development of multi-cellularity, so it is not surprising that such closely related isoforms would regulate similar cellular processes. Localization of FAs is tightly controlled by migrating cells, with coordinated assembly and turnover of FAs at the front of the cell body and release of adhesions at the rear. In Rab40b-4A and Rab40-3KO cells FAs are present throughout the cell and do not appear to be enriched specifically at leading lamellipodia, which helps explain difficulties in maintaining cell directionality.

Signaling through FAK is complex but recent reports have demonstrated its importance for regulating cell invasion. We observe differences in pFAK signaling in Flag-Rab40b-4A cells, which have greater levels of Y397 and less S910 compared to control and Flag-Rab40b cells. This result is in line with observations that phosphorylation of FAK at Y397 stabilizes FAs and decreases both invadopodia and ECM degradation, and that phosphorylation at S910 is associated with FA turnover, ECM degradation, and metastasis. While initially required to relay extracellular signaling cues, FAs must disassemble not only to generate the force needed for cell migration, but to free up signaling components required for invadopodia function [10, 11]. Taken together, our data demonstrate that the Rab40b/Cullin5 complex regulates cell migration and invadopodia formation, in part, by modulation of FA dynamics and FAK signaling, and are consistent with a model wherein Rab40b is central to regulating various aspects of cell migration and invasion.

The main function of SOCS box containing proteins is to serve as an adaptor to Cullin5 to mediate ubiquitylation of specific substrate proteins. Identification of these substrates is a key step to understand how Rab40b/Cullin5 mediates cell migration and invasion. Western Blot analysis of known CRL5 substrates showed no differences in stabilization in Rab40b-4A cells and suggests that the Rab40b/Cullin5 complex acts to regulate a unique set of proteins. In an effort to identify possible substrates, we completed comparative proteomic analysis of proteins that bind to either to wild type Flag-Rab40b or Flag-Rab40b-4A. We speculated that a subset of Rab40b bound substrates would not get ubiquitylated in the absence of CRL5 binding, and thus remain bound to Rab40b. Consistent with this idea, we identified several proteins that were enriched in mutant Flag-Rab40b-4A immunoprecipitate compared to wild-type. In this study we focused on EPLIN, an actin bundling protein and a well-established tumor suppressor [76]. Interestingly, EPLIN also binds to Rab40b in a GTP-dependent fashion. This raises an intriguing possibility that EPLIN act as a canonical Rab40b effector protein (Figure 10).

**Figure 10.**
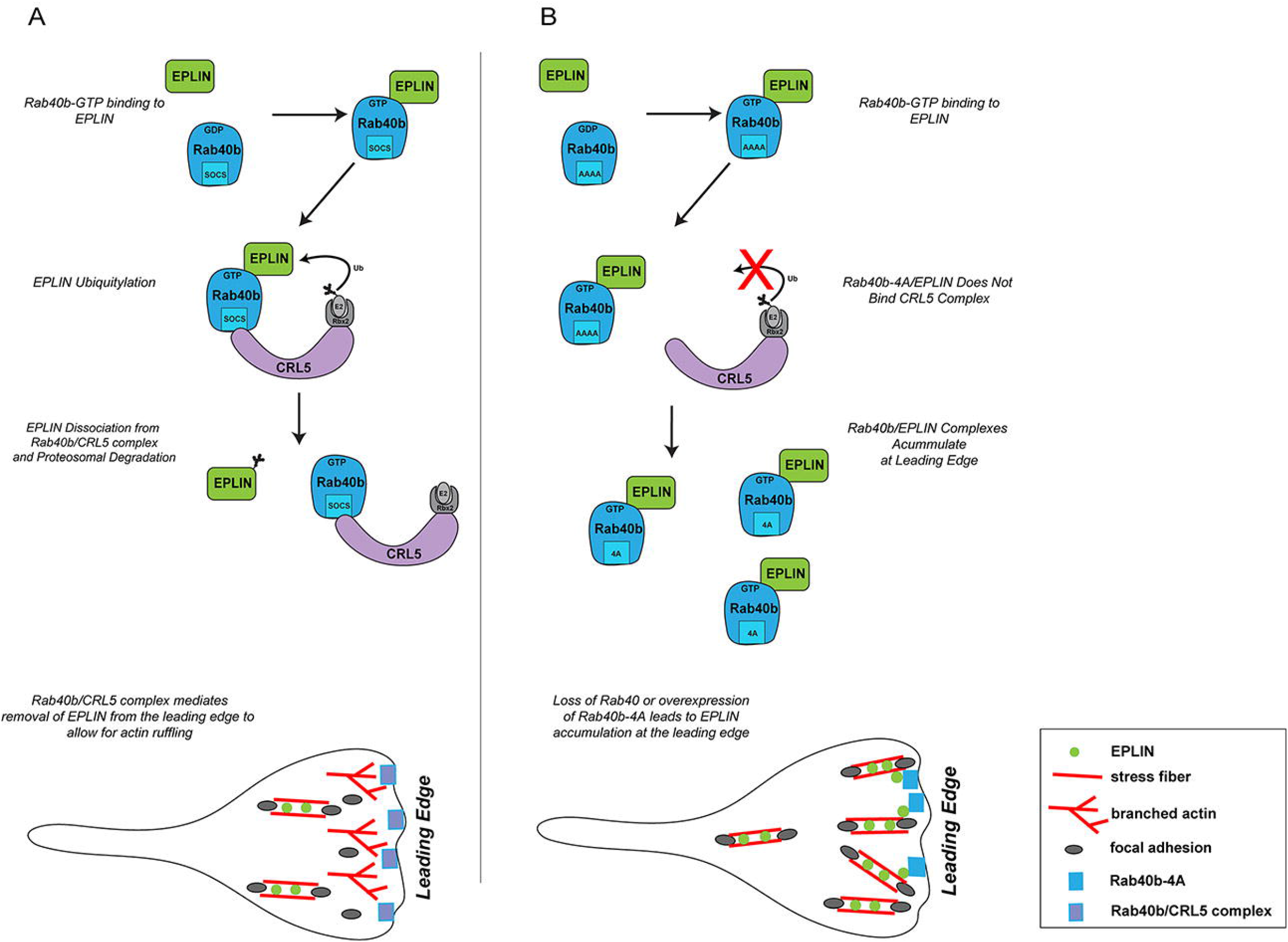
Proposed model for Rab40b function during cell migration. (A) GTP-bound Rab40b binds to EPLIN. Rab40b/EPLIN is then recognized by CRL5 complex via the Rab40b SOCS box. Binding of EPLIN with the Rab40b/CRL5 complex leads to EPLIN ubiquitylation, disassociation of EPLIN of Rab40b and EPLIN complex and eventual degradation by the proteasome. Since Rab40b is enriched at the leading edge of lamellipodia that leads to the exclusion of EPLIN from lamellipodia, thus, allowing actin ruffling. (B) Rab40b-4A mutation blocks Rab40b association with CRL5 complex, leading to inhibition of EPLIN ubiquitylation and stabilization of Rab40b-4A/EPLIN complex. Consequently, EPLIN accumulates at the leading edge of the lamellipodia, thus, resulting in inhibition of actin ruffling and increase in actomyosin stress fibers.

While it has been reported that EPLIN turnover occurs in response to growth factor stimulation [77], the molecular machinery that facilitate this process is not known. Here we show that Rab40b binds to Cullin5 and that loss of Rab40b decreases levels of ubiquitin-conjugated EPLIN, thus leading to a decrease in EPLIN degradation. This suggests a model in which EPLIN binds Rab40b, which in turn binds Cullin5 and the remaining components of the CRL5, leading to ubiquitylation of EPLIN (Figure 10A). The EPLIN ubiquitylation then leads to dissociation of Rab40b-EPLIN complex and subsequent EPLIN degradation (Figure 10A). Although an upstream signaling mechanisms that triggers EPLIN ubiquitylation have been reported, to our knowledge we are the first to identify specific molecular components responsible for EPLIN ubiquitylation and degradation. In addition to proteosomal degradation, cells utilize ubiquitylation of target proteins to regulate activity and localization and we observe that Rab40b/Cullin5 alters the localization of EPLIN to the leading edge of lamellipodia, presumably by localized EPLIN ubiquitylation. Indeed, the addition of ubiquitin as a standard post-translational modification has been shown to regulate localization and activity of some target proteins [78, 79]. Interestingly, we could only detect Rab40-dependet ubiquitylation of EPLIN-α isoform. Although we have not directly addressed the mechanism that determines the discrepancy in regulatory fates of EPLIN isoforms, it is possible that our assays were simply not sensitive enough to detect transient ubiquitylation in EPLIN-β. Alternatively, since it has been suggested that EPLIN-α expression regulate EPLIN-β, it could be that EPLIN-α ubiquitylation also indirectly affects EPLIN-β function and localization.

The high sequence similarity amongst the Rab40 family members makes it likely that other Rab40 isoforms may play a role in EPLIN binding, stabilization or localization. Indeed, the fact that Rab40b/Rab40c double knock-down further decreased EPLIN-α ubiquitylation (as compared to Rab40b knock-down) suggests that the entire Rab40 family may be involved in regulating EPLIN. Consistent with this hypothesis, only cells lacking Rab40a, -b and -c isoforms phenocopy defects observed in cells expressing Flag-Rab40b-4A. The Flag-Rab40b-4A mutant stabilizes the Rab40b and EPLIN interaction, and its expression likely leads to formation of Rab40b/EPLIN complexes that are unrecognized by CRL5 components. The accumulation of such Rab40b complexes with EPLIN or other substrates makes it likely that mutant Rab40b-4A acts in a dominant-negative fashion with respect to Cullin5 binding. Ascertaining substrate specificity and overlap for different Rab40 isoforms, as well as how Rab40/substrate interactions impact cellular function, will be the focus of our future work.

Here we show that Rab40b localizes to lamellipodia, where it colocalizes with actin ruffles, independent of Cullin5 binding capability, and that these actin ruffles are diminished upon expression Rab40b-4A or depletion of Rab40a/b/c. Coincident with a decrease in lamellipodia actin ruffles, we observe an accumulation of EPLIN at the leading edge and a decrease in lamellipodia dynamics. The ability of EPLIN to bundle actin filaments and decrease actin dynamics, together with its regulation by Rab40b/Cullin5, suggests to us that it is likely Rab40b/Cullin5-dependent ubiquitylation is needed to remove and exclude EPLIN form the leading edge for efficient cell migration to occur. Consistent with this hypothesis, overexpression of Rab40b-4A or knockout of Rab40a/b/c leads to accumulation of EPLIN at the leading edge, thus, leading to a decrease in leading edge dynamics. We propose that rather than regulating global cellular EPLIN levels, Rab40b regulates EPLIN function via localized ubiquitylation-dependent removal and degradation from the leading edge of the cell (Figure 10).

In contrast to previous reporting [70], overexpression of either EPLIN isoform was not sufficient to drive cells to a prominent stress fiber phenotype in our system. In addition to inherent cell-type specific differences, this suggests that other molecular factors in addition to EPLIN contribute to the observed phenotypes in Rab40b-4A mutant or Rab40-3KO cells. Interestingly, the majority of proteins that exhibit increased binding to Rab40b-4A are regulators of actin cytoskeleton dynamics, suggesting that the Rab40b/Cullin5 complex targets a specific subset of actin regulators to effect actin dynamics during cell migration (Figure 5A). For example, ARHGAP42 [80] and ARHGAP19 [81] both regulate RhoA activation. LMO7 is a LIM domain containing protein implicated in the regulation of actin dynamics at FAs and adherens junctions [82]. Finally, DIAPH1 is an actin nucleating factor that regulates actin polymerization in response to RhoA activation [83]. While further studies will be needed to confirm and identify specific roles of these Rab40b binding proteins, the common feature amongst them all is that they are involved in the regulation of RhoA signaling and actin dynamics.

In this study, we also analyzed the role of Rab40b/Cullin5 binding in tumor growth *in vivo* using an orthotopic xenograft model. We found that loss of Rab40b/Cullin5 binding resulted in a reduction of tumor growth. Our previous work has demonstrated loss of Rab40b-mediated MMP2/9 secretion leads to decreased growth potential [18] since MMP release both remodels the ECM and releases insoluble cytokines which stimulate angiogenesis, cell migration and proliferation. In addition to defects in MMP2/9 secretion, loss of proper chemotaxis response has been demonstrated to inhibit *in vivo* growth, thus it is likely that the Rab40b/Cullin5 effects are also applicable in a 3D context. There is a similarly well-established role for FAK signaling in the promotion of cancer. In addition to regulating focal adhesion dynamics in motile cells, it is likely that altered FAK signaling contributes to a reduction in tumor growth in Rab40b-4A cells. In line with cell culture models, *in vivo* tumors saw increased EPLIN staining, which has an established role in regulating tumor growth. This suggests, that despite some differences between tissue culture and *in vivo* systems, EPLIN is still regulated by a Rab40b/Cullin5 mechanism. Surprisingly, we also see increased E-cadherin staining in xenograft tumors, particularly along the tumor border with the tumor microenvironment. While E-cadherin is required for 2D collective movement in some contexts, it is known to inhibit migration in an *in vivo* context. Hence, the upregulation of E-cadherin at the boundary of the tumor with the extracellular space makes it interesting to speculate that the inability of cells to properly interact with their external environment causes a partial reversion from an invasive mesenchymal-like state to a more epithelial, sedentary state. This alongside our previous data showing that Rab40b regulates MMP secretion suggest that multiple aspects of tumor invasion/migration are regulated by Rab40b.

Taken together, our data suggest that Rab40b is a unique small monomeric GTPases that appears to have two distinct functions. On one hand, Rab40b functions as a canonical Rab GTPase by regulating MMP2/9-containing secretory vesicle targeting to invadopodia [14–16]. On the other hand, Rab40b acts as a SOCS-adaptor protein for CRL5 and mediates Cullin5-dependent protein degradation. These two functions seem to converge on the same downstream goal to regulate subcellular localization of specific actin regulators and to promote directional cell migration and invasion. Many questions, however, still remain. How is Rab40b and EPLIN binding regulated? Does Rab40b regulate cell migration during development *in vivo?* How rab40b is targeted to the leading edge of the lamellipodia? What are the functions of other Rab40 isoforms and substrates? Additional work will be needed to address these questions and will be the focus of future studies.

## MATERIALS AND METHODS

### Cell Culture and Cell Lines

All cell lines were cultured as described previously ([17]). MDA-MB-231 cell line stably expressing Flag–Rab40b-4A was created by cloning Rab40b-4A (primers purchased from IDT, Coralville, IA) into lentiviral pCS2-FLAG vector obtained from Addgene (Cambridge, MA). Cell lines were routinely tested for mycoplasma. All cell lines used in this study were authenticated and are in accordance with ATCC standards. For all Flag-Rab40b and Flag-Rab40b-4A experiments we used parental MDA-MB-231 cells as controls. For all Rab40-3KO experiments MDA-MB-231 cells expressing tet-inducible Cas9 were used as a control.

### Plasmids

GFP-Paxillin, GFP-EPLIN-α, and GFP-EPLIN-β-were purchased from Addgene (50529, 40948, 40947). For Flag-Rab40b-4A mutants were generated from an existing PLVX-FLAG-Rab40b plasmid [17]. Point mutations for 213L and 215T were created by site directed mutagenesis with the following primers:

213L Forward (5’ GTGGACAAGCTCCTGCTCCCCATTGCC 3’);

213L Reverse (5’ GGGAATGGGGAGCAGCTTGTCCAC 3’);

215T Forward (5’ GACAAGCTCCCGCTCACCATTGCCTAA 3’);

215T Reverse (5’ CTTAAGGCAATGGTGAGCGGGAGCTTG 3’).

For the Flag-Rab40b-4A mutation, the entire coding sequence for Rab40b containing the AAAA mutation (CTCCCGCTCCCC –> GCAGCAGCA) was ordered from Integrated DNA Technologies, Inc. (Carolville, IA) on a Pbluescript plasmid and subcloned into an existing PLVX-FLAG-Rab40b plasmid. Lentiviral transfection of Rab40b mutant plasmids into MDA-MB-231 cells was then performed as previously described [17].

### Antibodies

Antibodies used in this study are as follows: anti-Flag – (WB 1:1000, Sigma F3165), anti-EPLIN – (WB 1:1000, IF 1:100, Cell Signaling Technology 50311), anti-EPLIN (WB 1:1000, IF 1:100, Sant Cruz Biotechnology sc-136399), anti-EPLIN (WB 1:1000, IP 1 ug/1 mg lysate, IHC 1:200 Proteintech 16639-1-AP), anti-HA (WB 1:500, IP 2 μg/1mg cell lysate, Santa Cruz SC F-7), Alexa-Fluor-568–phalloidin (IF 1:1000, Life Technologies A1238), SiR-Actin – (1:10,000, Cytoskeleton CY-SC001), anti-β-tubulin – (WB 1:2500, LiCor 926-42211), anti-α-tubulin – (WB 1:2500, Santa Cruz 23948), pFAK Y397 (WB 1:1000, Abcam ab8129), FAK S910 (WB 1:1000, Invitrogen 44-596G), anti-E-Cadherin (WB 1:1000, IHC 1:200, Cell Signaling 3195), total FAK (WB 1:1000, BD Biosciences 610087), anti-Zyxin (IF 1:1000 abcam, ab50391), anti-Rab40b (WB 1:500, LSBio, LS-C353287), anti-Rab40c (WB 1:500 Santa Cruz, H-8 sc-514826), anti-CD63 (IF 1:100, gift from Dr. Andrew Peden).

### 3D Inverse Invasion Assay

Assay were adapted from von Thun et al., 2012 and performed as previously described. In brief, a Matrigel (BD Biosciences Bedford, MA) plug supplemented with 50 μg/ml fibronectin was made on a transwell filter (Corning Life Sciences Tewksbury, Massachusetts, Cat. #3422). The cells were allowed to invade towards a gradient of 10% fetal bovine serum (FBS) and 10% Nu serum for 7 days. The cells were stained with 4 μM Calcein for 15 min and imaged at 5-μm steps to a total distance of 120 μm. ImageJ software was used to quantify the number of cells in every 5-μm step image from 5 μm to 100 μm. Images were analyzed as previously described [18]. For quantification, at least 20 cells from three different fields per treatment were counted.

### Migration Assays and Time-Lapse Analyses

For scratch assays, wells of a 96-well dish were coated with rat tail collagen (Corning Life Sciences Tewksbury, Massachusetts, Cat. #354249) and allowed to set for 1 hours at room temperature. Wells were then washed once with PBS and cells were plated to produce a confluent monolayer 24 hours later and serum starved overnight. A WoundMaker™ (Essen BioScience, Ann Arbor, Michigan) was used to make scratches across each well. Wells were then rinsed twice with PBS and plated in low-serum media (DMEM + 2% FBS). Images were acquired over the center of the scratch every 2 hours for 48 hours. IncuCyteZOOM software was used to measure percent wound closure over time.

Chemotaxis assays were performed with the IncuCyte^®^ Live-Cell Analysis System according to manufacturer protocol. In brief, 1000 sub-confluent, serum starved cells were plated into either fibronectin coated or collagen-coated upper wells of a 96 well plate (IncuCyte^®^ ClearView 96-well Chemotaxis Plate, cat # 458). Upper chambers were supplied either regular growth serum (DMEM + 10%FBS) or low serum (2% FBS). The top and bottom wells were imaged every two hours for 48 hours. IncuCyte^®^ analysis software was used to analyze and quantify changes in cell area over time, normalized to the initial top chamber seeded cell count.

For individual cell time-lapse migration analysis, cells were plated on collagen-coated 35mm glass-bottomed dishes. Cells were serum starved overnight in media supplemented with SiR-actin. Cells were then washed once with PBS supplemented with DAPI at 1:20,000, once again with PBS and then placed in full growth media supplemented with SiRactin. Cell dishes where then placed in a climate controlled chamber and images every 10 minutes. Images were taken on either a Zeiss LSM 880with Zen Blue software (ZEISS, Oberkochen, Germany) with or a Nikon A1R confocal system with NIS Elements software (Nikon, Tokyo, Japan) in a live imaging chamber. Individual cell migration dynamics for individual cells was analyzed using ImageJ and the Manual Tracking plug-in (Fabrice Cordelires, Institut Curie, Orsay, France). Generated values were then used to determine mean-squared displacement, velocity and directionality as described in [84].

For time-lapse analysis of focal adhesion sites, cells were transfected with GFP-Paxillin with jetPRIME (Polyplus, Berkeley, CA) according to manufactures recommendations. Cells were then seeded onto a collagen coated glass-bottom dish and incubated overnight with SiR-actin in minimal media. One hour prior to imaging, cells were washed once with PBS and incubated in full growth media supplemented with SiR-actin. Cell dishes where then placed in a climate controlled chamber and imaged every 4 minutes on Zeiss LSM 880. Images were pre-processed in ImageJ and uploaded to the FAAS server (https://faas.bme.unc.edu/) [85] to measure adhesion dynamic characteristics. Adhesions tracked less than three consecutive frames were discarded. Lifespan was determined only for adhesions whose entire lifetime occurred during imaging.

To determine lamellipodia dynamics MDA-MB-231 cells (control, Flag-Rab40-3KO and Flag-Rab40b-4A) were plated on collagen-coated glass-bottom dish and incubated overnight. Cell dishes were then placed in a climate-controlled chamber at 37C° and imaged by DIC using time-lapse microscopy. In all cases, 50 images with time lapse of 3 seconds were taken for each cell. Images were taken on either a Zeiss LSM 880with Zen Blue software (ZEISS, Oberkochen, Germany). To determine the lamellipodia dynamics, the single pixel was chosen on the edge of the lamellipodia and DIC signal was measured at every time point. The max-intensity of DIC signal was than calculated and plotted along X-axis.

### Mouse mammary fat pad xenograft assays

At total of thirty 8-week-old female hairless athymic nude mice were divided into three experimental groups with 10 mice per group. Two million log phase MDA-MB-231 cells (control, stably expressing wild-type Flag-Rab40b, or stably expressing Flag-Rab40b-4A) were injected into the number four right and left intact mammary glands. Primary tumor growth was measured twice weekly using calipers. Once total tumor burden was reached (2 cm^3^ for a single tumor or 3 cm^3^ total tumor burden), or tumors had exhibited ulcerations, or moribound criteria were met the mice were humanely euthanized. Mammary tumors and lungs were harvested from each animal. One half of the tissues were flash frozen for further analysis. The other half were formalin fixed and paraffin embedded for histology and IHC analysis. All animal experiments were performed according to IUCAC approved guidelines.

### Quantitative immunohistochemical analysis

Mammary glands (tumors intact) were harvested and placed into 10% neutral buffered formalin for 48 h. After 48 h, tissues were moved to 70% EtOH, processed, embedded, sectioned and stained for H&E or IHC as described [86–88]. Immunofluorescent images were obtained using 40X magnification on OLYMPUS microscope for analysis. For Ki67, EPLIN and E-Cadherin, stain quantification of total tumor area (necrotic and stromal areas removed) and percent positive stain or stain intensity was performed using ImageScope Aperio Analysis software (Leica, Buffalo Grove, IL). Areas for quantification were annotated using Aperio analysis tools, and percentage of weak, medium, and strong pixels was determined using the color deconvolution algorithm.

### Flag–Rab40b immunoprecipitation and proteomic analyses

Putative Rab40b-binding proteins were identified by co-immunoprecipitation using anti-Flag antibody coated beads as described previously [89]. Proteins eluted from anti-Flag antibody beads were then analyzed using tandem mass spectrometry analysis as follows. Protein samples were digested with trypsin using the Filter-Aided Sample preparation (FASP) protocol [86]. Liquid chromatographic analysis was performed in a Waters Acquity ultra performance LC system (Waters Corporation, Wilmslow, UK). Peptide separation was performed on an ACQUITY UPLC HSS T3 250 mm analytical column. Data were acquired using Synapt G2 HDMS mass spectrometer and Masslynx 4.1 software (Waters Corporation) in positive ion mode using data independent (DIA) acquisition Raw data were lock mass-corrected using the doubly charged ion of [Glu1]-fibrinopeptide B (*m/z* 785.8426; [M+2H]2+).

Raw data files were processed and searched using ProteinLynx Global SERVER (PLGS) version 3.0.1 (Waters Corporation, UK). Data was analyzed using trypsin as the cleavage protease, one missed cleavage was allowed and fixed modification was set to carbamidomethylation of cysteines, variable modification was set to oxidation of methionine. Minimum identification criteria included 2 fragment ions per peptide, 5 fragment ions per protein and minimum of 2 peptides per protein. The following parameters were used to generate peak lists: (i) minimum intensity for precursors was set to 135 counts, (ii) minimum intensity for fragment ions was set to 25 counts, (iii) intensity was set to 750 counts. UniprotKB/SwissProt human database was used for protein identification.

All candidate proteins identified in mass spectrometry were then filtered using two following criteria: 1) candidate proteins had to be enriched at least 10 fold over IgG control, 2) all RNA, DNA-binding proteins, as well as mitochondria proteins were eliminated as putative contaminants. Finally, only proteins enriched more than 2 fold in Flag-Rab40b-4A samples were considered as putative Rab40b/Cullin5 substrate proteins (Figure 1B). Full list can be found in Supplemental Table 1.

### Flag-Rab40b-4A proteomics

In brief, wild-type Flag-Rab40b or Flag-Rab40b-4A were N-terminally Flag-tagged, expressed ubiquitously in MDA-MB-231 breast cancer cells, and immunoprecipitated using Flag beads prior to mass spectrometry analysis. To determine Rab40b-interacting proteins (both WT and 4A), we established the following criteria. First, only proteins >3-fold IgG control (spectral counts) were analyzed. Second, any hits identified as nonspecific based on the CRAPome database were dismissed, as well as any additional DNA, RNA, and mitochondrial proteins. Finally, a 2-fold enrichment cutoff (spectral counts) was used to identify proteins preferentially bound to Flag-Rab40b-4A vs Flag-Rab40b. After analyzing protein hits from two independent experiments, we identified a set of Rab40b binding proteins that were enriched in Flag-Rab40b-4A compared to wild-type Rab40b. From the first run, 43.1% (81/188) of proteins were enriched in 4A vs WT. In the second run, 52.5% (32/61) of proteins were enriched in 4A vs WT. Proteins listed in Figure 5A are a shortened list of putative substrates, highlighting the theme of actin regulators. Full list can be found in Supplemental Table 1.

### Reverse transcriptase polymerase chain reaction (RT-PCR) and quantitative PCR (qPCR)

Total RNA was extracted from 2×10^7^ MDA-MB-231 cells using TRIzol (Invitrogen) according to the manufacturer’s protocol. Reverse transcription to cDNA was performed with SuperScript III (Invitrogen) using random hexamer primers. PCR was performed using Taq polymerase (Invitrogen). To quantify the percent of knockdown, cDNA from mock-or siRNA-treated cells was analyzed in triplicate by qPCR amplification using Sybr Green qPCR Master Mix using Applied Biosystems ViiA7 Real-Time PCR System. The qPCR amplification conditions were: 50°C (2 minutes), 95°C (10 minutes), 40 cycles at 95°C (15 seconds), 60°C (1 minute). Primer pairs were designed to amplify mRNA-specific fragments and unique products were tested by melt-curve analysis. Amplification efficiency was calculated using the slope of the regression line in the standard curve. Targets were normalized to GAPDH. The primers used for qPCR are from PrimerBank (https://pga.mgh.harvard.edu/primerbank/) and are listed as follows:

RAB40A Forward (5’-CTGCGGCACAGGATGAATTG-3’)

RAB40A-Reverse (5’-AGGCTGCTCTTGTGAGTGGA-3’)

RAB40B- Forward (5’-GTCCGGGCCTACGACTTTC-3’)

RAB40B- Reverse (5-GGCCTGAAGTATCCCAGAGC-3’)

RAB40C- Forward (5’-GGCCCAACCGAGTGTTCAG-3’)

RAB40C- Reverse (5’-GGACTTGGACCTCTTGAGGC-3’)

### Zymography assay

Control, Flag-Rab40b and Flag-Rab40b-4A MDA-MB-231 cells were incubated in the complete medium at 37°C. After 24 h of incubation, the medium was replaced with Opti-MEM (Invitrogen, Carlsbad, CA) and cells were incubated at 37°C for another 24 h. Cell medium was collected and briefly centrifuged to clear the lysate. Equal concentrations of each sample were loaded into a 7.5% acrylamide gel supplemented with gelatin type A from porcine skin. After running the gel, gels were washed with water and incubated twice in 2.5% TritonX-100 for 30 minutes. Gels were then incubated in substrate buffer 50mM Tris, pH8.0, 5mM CaCl_2_) for 48 hours at 37°C. Gels were stained with Coomassie for 1 hour, de-stained and imaged on Gel Doc™ EZ Imager (Bio-Rad Laboratories, Hercules, CA). MMP2/9 secretion was quantified using Quantity One Software (Bio-Rad).

### siRNA Knockdown

For EPLIN, siRNAs were purchased from Dharmacon (Dharmacon, Lafayette, CO). D001210-01-05 (non-targeting), D010663-02-0002 (LIMA1 #1), and D010663-03-0002 (LIMA1 #2) siRNAs were transfected using JetPrime according to manufacturer protocol. Cells were trypsinized and re-plated onto collagen coated coverslips at 24 hours. Cells were fixed at 48 hours, stained and imaged and the A1R. For Rab40b and Rab40c, siRNAs were purchased from Sigma. Mission siRNA universal negative control (SIC001, sigma), Rab40b siRNA (SASI_Hs01_00150368) and Rab40c siRNA (SASI_Hs01_00031938) were transfected using lipofectamine RNAiMAX (Invitrogen) according to manufacturer protocol.

### Immunofluorescence microscopy

MDA-MB-231 cells were seeded onto collagen-coated glass coverslips and grown in full growth media unless otherwise noted for at least 24 hours. Cells were washed with PBS and fixed in 4% paraformaldehyde for 15 minutes. Samples were then washed 3x in PBS then incubated in blocking serum (1X PBS, 5% normal donkey serum, 0.3% Triton X-100) for 1 hour at room temperature. Primary antibodies were then diluted at 1:100 in dilution buffer (1X PBS, 1% BSA, 0.3% Triton X-100) overnight at 4°C. Samples were then washed 3x with PBS, then incubated with fluorophore-conjugated secondary antibodies (1:100 in dilution buffer) for 1 hour at room temperature. Cells were then washed 3x in PBS and mounted onto glass slides. Cells were then imaged on either an inverted Zeiss Axiovert 200M deconvolution microscope with a 63× oil-immersion lens and Sensicam QE CCD camera or a Nikon A1R. Z-stack images were taken at a step size of 100–500 nm.

### CRISPR/Cas9 Knock-out lines

Rab40 CRISPR 3KO cell lines were generated using MDA-MB-231 cell line stably expressing tet-inducible Cas9. For Rab40a, guide RNAs targeting CCTCAAAGAAGGTCACACCC (Exon 3) and TTGGCTCGGGAGGCCGAGCA (Exon 3) were transfected into Dox-induced Cas9 MDA-MB-231s. For Rab40b, guide RNAs targeting TCCAGGGATACTTCAGGCCA (Exon 3) and TCTGGCGGCCGAGCAAGGGT (Exon 5) were transfected into Dox-induced Cas9 MDA-MB-231s. For Rab40c, guide RNAs targeting TACCGTTACTGTAGGCGTAC (Exon 1) and AGGTAGTCGTAGCTCTTCAC (Exon 3) were transfected into Dox-induced Cas9 MDA-MB-231s. Cells were split 24 hours after transfection and seeded for single colonies. Clones were screened by PCR for followed by TA cloning and genotyping. First, Rab40b and Rab40c double KO (2KO) was generated. One of the Rab40b/c 2KO clones were than used to generate Rab40a/b/c 3KO. Two different 3KO lines were then used for experiments. The mutations in all Rab40a, Rab40b Rad rab40c alleles are listed below (asterisk marks where mutant allele sequence diverges from wild type due to frame shift mutation):

Rab40c allele (both alleles have the same mutations):
MGSQGSPVKSYDYLLKFLLVGDSDVGKGEILESLQDGAAESPP*TVTGSTTRPPPSCWTAGA

Rab40b allele #1:
MSALGSPVRAYDFLLKFLLVGDSDVGKGGEILASLQDGAAESPYGHPAGIDYKTTTILLDGRRVKLQL WDTS*AREDFVPYSAPTPGAHRV

Rab40b allele #2:
MSALGSPVRAYDFLLKFLLVGDSDVGKGGEILASLQDGAAESPYGHPAGIDYKTTTILLDGRRVKLQLWDTSGQGRFCTIFRSYSRGAQGVILVYDIANRWSFDGIDRWIKEIDEHAPGVPKILVGNRLHLAFKRQV PTEQAQAYAERLGVTFFEVSPLCNFNITESFTELARIVLLRHGMDRLWRPSK*C

Rab40C allele #1 (Rab40-3KO-1 line):
MSAPGSPDQAYDFLLKFLLVGDRDVGKSEILESLQDGAAESPYSHLGGIDYKTTTILLDGQRVKLKLWDTSGQGRFCTIFRSYSRGAQGVILVYDIANRWSFEGMDRWIKKIEEHAPGVPKILVGNRLHLAFKRQVP REQAQAYAERLGVTFFEVSPLCN*FNIIESFTELARIVLLRHRMNWLRPSKVLSLQDLCCRTIVSCTPVH LVDKLPLPSTLRSHLKSFSMAKGLNARMMRGLSYSLTTSSTHKSSLCKVEIVCPPQSPPKNCTRNSCKIS

Rab40C allele #2 (Rab40-3KO-1 line):
MSAPGSPDQAYDFLLKFLLVGDRDVGKSEILESLQDGAAESPYSHLGGIDYKTTTILLDGQRVKLKLWDTSGQGRFCTIFRSYSRGAQGVILVYDIANRWSFEGMDRWIKKIEEHAPGVPKILVGNRLHLAFKRQVP REQAQAYAERLG*GGRARY

Rab40C allele #1 (Rab40-3KO-2 line):
MSAPGSPDQAYDFLLKFLLVGDRDVGKSEILESLQDGAAESPYSHLGGIDYKTTTILLDGQRVKLKLWDTSGQGRFCTIFRSYSRGAQGVILVYDIANRWSFEGMDRWIKKIEEHAPGVPKILVGNRLHLAFKRQVP REQAQAYAERL*GVTFFEVSPLCNFNIIESFTELARIVLLRHRMNWLRPSKVLSLQDLCCRTIVSCTPVH LVDKLPLPSTLRSHLKSFSMAKGLNARMMRGLSYSLTTSSTHKSSLCKVEIVCPPQSPPKNCTRNSCKIS

Rab40C allele #2 (Rab40-3KO-2 line):
MSAPGSPDQAYDFLLKFLLVGDRDVGKSEILESLQDGAAESPYSHLGGIDYKTTTILLDGQRVKLKLWDTSGQGRFCTIFRSYSRGAQGVILVYDIANRWSFEGMDRWIKKIEEHAPGVPKILVGNRLHLAFKRQVP REQAQAYAERL*premature STOP codon

### Glutathione bead pull-down and immune-precipitation assays

Glutathione bead pull-down assays were performed as previously [90]. Briefly, glutathione beads (GoldBio St Louis, MO) were coated with GST-Cullin5 or GST control in PBS and incubated with MDA-MB-231 control or Rab40b-Mutant cell lysates (PBS, 1% Triton X-100, 10mM PMSF) in a final volume of 0.5□ml of reaction buffer (50□mM HEPES, pH 7.4, 150□mM NaCl, 0.1% Triton X-100, and 1□mM PMSF). Samples were incubated at 25□°C for 1□h while rotating, pelleted by centrifugation at 2,000*g* for 3□min, rinsed with reaction buffer (1□ml × 3), and bound proteins were eluted with 1% SDS, analyzed by SDS–PAGE, and stained with Coomassie blue or immunoblotted and scanned using a LI-COR Odyssey scanner (LI-COR Biosciences, Lincoln, NE).

For EPLIN and FLAG-Rab40b co-immunoprecipitation assays, 1 ml of a 1mg/ml of cell lysates were pre-cleared with 25 ul of Protein G sepharose beads for 1 hour at 4°C, then incubated with 5 μg of IgG control (115-006-006, Jackson Laboratory, Bar Harbor, ME) or anti-FLAG antibody (Protein Tech 16639-1-AP) and incubated at 37°C while rotating. After 60min, 50 μl of Protein G beads were added and samples were incubated at 37°C for 60 min while rotating. Samples were then washed pelleted by centrifugation at 2,000*g* for 3 min then washed 4x in wash buffer (50□mM HEPES, pH 7.4, 300□mM NaCl, 0.1% Triton X-100, 0.1% SDS). Bound proteins were eluted with 1% SDS, analyzed by SDS–PAGE, immunoblotted and scanned using a Li-Core Odyssey scanner.

### EPLIN Ubiquitylation Assay

293t cells were plated into 10cm dishes. When cells were 50% confluent, cells were transfected with siRNAs listed above (control, Rab40b, Rab40c, or both). After 24 hours, were transfected with FLAG-α-EPLIN using Lipofectamine 2000 (Invitrogen). After 24hrs, cells were treated with 10μM MG132 overnight then lysed in Tris-buffer containing 10mM Iodoacetamide (IAA). Cell lysates were subject to IP with FLAG antibody (M2, Sigma) followed by Western Blot with FLAG or Ubiquitin antibody (P4D1, Santa Cruz).

### Image analysis

#### Analysis of Focal Adhesion number and size

Focal adhesion quantification performed in ImageJ and was adapted from [91]. In brief, maximum intensity projections for relevant z-planes were created and images were loaded into ImageJ. Background was minimized using the Subtract Background and EXP tools, images were filtered using the Log3D/Mexican Hat plugin, thresholds were then applied manually (method = default). Individual cells were defined by hand, and focal adhesions were determined with the Analyze Particle command. Resulting particle outlines were then compared to the original image to ensure fidelity of the analysis. For live imaging analysis of focal adhesion, images were processed in ImageJ and uploaded to the FAAS server [85].

#### Analysis of FAs positive for zyxin

To determine colocalization of zyxin positive focal adhesions, zyxin/paxillin dual stained images were first split using the Split Channels tool in ImageJ, background was subtracted and thresholds were adjusted as above. Individual cells were individually defined manually, then analyzed with the Colocalization plugin (Pierre Bourdoncle, Institut Jacques Monod, Service Imagerie, Paris) to determine zyxin-positive paxillin patches. Determination for cortactin-positive actin puncta was done similar to that of zyxin-positive focal adhesions.

#### Quantification of cellular EPLIN and non-muscle Myosin IIA/B

To measure the levels of EPLIN and non-muscle Myosin IIA/B associated with actin cytoskeleton, MDA-MB-231 cells (control, Rab40-3KO and Flag-Rab40b-4A) were fixed and stained with phalloindin-Alexa594 and either EPLIN or non-muscle Myosin IIA/B antibodies. Randomly selected cells from two random fields were then imaged using the same exposure time. Masks where than generated for each cell and sum-fluorescence intensity of EPLIN or non-muscle Myosin IIA/B was calculated for each cell. The sum-intensity was then expressed as fluorescence per 1 μm^2^ for each cell analyzed. The experiment was repeated three times and the values expressed as means and standard deviations calculated from all cells analyzed.

#### Cell spreading analysis

To determine the effect of Rab40b on cell-ECM adhesion, MDA-MB-231 cells (control, Rab40-3KO and Flag-Rab40b-4A) were plated on collagen-coated coverslips and incubated for 90 min. Cells were then fixed and stained with phalloindin-Alexa594. Randomly selected cells from three random fields were then imaged. Masks where than generated for each cell and the area covered by each individual cell was than calculated. The experiment was repeated three times and the values expressed as means and standard deviations calculated from all cells analyzed.

#### Line-scan analysis of EPLIN distribution in lamellipodia

To determine the subcellular distribution of EPLIN, MDA-MB-231 cells (control, Rab40-3KO and Flag-Rab40b-4A) were plated on collagen-coated coverslips and incubated for 24 hours. Cells were then fixed and stained with anti-EPLIN antibodies and phalloidin-Alexa594. For individual isoforms, cells were plated on collagen-coated coverslips and incubated overnight, then transfected with GFP-Eplin-α or HPLIN-β then incubated for 24 hours. Cells were then fixed and stained with anti-EPLIN antibodies and phalloidin-Alexa594. Z-stack images were acquired and maximum projection intensities were created of each image. Only cells with well-defined lamellipodia (as determined by imaging phalloidin-Alexa594) were chosen for this analysis. Lines were then drawn across the lamellipodia perpendicular to nucleus. The intensity of EPLIN and F-actin in each pixel along this line was determined with either ImageJ or 3i Slidebook imaging software. Intensities along each lines were normalized, and the front edge of either actin or EPLIN was determined were intensity was at least 20 percent of maximum.

### Statistical Analysis

Statistical analysis for all experiments was determined using GraphPad Prism Software (GraphPad, San Diego, CA). A two-tailed Student’s t-test was used to determine statistical significance unless otherwise noted. Data was collected from at least three independent experiments unless otherwise noted. In all cases p ≤ 0.05 was regarded as significant. Error bars represent standard erros unless otherwise noted. For all immunofluorescence experiments, at least five randomly chosen image fields per condition were used for data collection. For quantitative immunofluorescence analysis, the same exposure was used for all images in that experiment and quantified using ImageJ.

## ACKNOWLEDGEMENTS

We are grateful to the laboratory of Dr. Rebecca Schweppe for guidance on Focal Adhesion Kinase experiments. We also thank Veronica Wessells, Alan Elder and Sarah Tarullo from Dr. Traci Lyons laboratory for assistance with mouse work. We also appreciate the Section of Developmental Biology in the Department of Pediatrics and the Gates Frontiers Fund for microscopy support, as well as Tim Vanderleest at University of Denver for help with cell migration kinetics. HA-ubiquitin constructs were a gift from Changwei Liu. This work was funded by R01 GM122768 to RP, Bolie Family Foundation fellowship to EL, and T32 GM008730 to ED.

## FIGURE LEGENDS

**Supplemental Figure 1**

(A) Table listing other proteins (in addition to Figure 1) identified by mass spectroscopy analysis from Flag-Rab40b pulldown.

(B) Mean Squared Displacement calculated from time-lapse analysis of parental (control) and Flag-Rab40b-4A cells (also see Supplemental Movies 1 and 2). Data shown are the means and SEM derived from n=39 control cells and n=79 Flag-Rab40b-4A cells. p = 0.75. For power-law equation fitting to MSD(Δt) = C*Δt^α^, where the exponent α is indicative of type of movement, control - MSD(Δt) = 0.8337Δt^1,3212^, α= 1.3212; Flag-Rab40b-4A - MSD(Δt) = 0.6652Δt^1,3743^, α = 1.3743.

(C) Average speed calculated from time-lapse analysis of parental (control) and Flag-Rab40b-4A cells (also see Supplemental Movies 1 and 2). Control - 0.24 μm per minute ± 0.0 SEM, n=51 cells; Flag-Rab40b-4A - 0.25 μm per minute ± 0.01 SEM, n=121.

(D) Directionality ratios control and Flag-Rab40b-4A cells were calculated from time lapse analysis (also see Supplemental Movies 1 and 2). Only cells that remained in frame for the duration of the entire time-lapse experiment were analyzed. Directionality at last time point for control cells is 0.38 ± 0.03 SEM and is 0.29 ± 0.02 SEM for Flag-Rab40b-4A cells. p < 0.05.

(F) Rates of cell proliferation of control and Flag-Rab40b-4A cells on either collagen or fibronectin. The data shown are the means and SEM derived from three different experiments.

**Supplemental Figure 2**

(A) Migration analysis of control, Flag-Rab40b and Flag-Rab40b-4A cells by scratch assay. Right; Representative images of cells at various time points. Blue boxes designate computer generated boundaries of original scratch border. Left; Quantification of three separate runs, at least 5 wells per condition per run.

(B) Images of control and Flag-Rab40b-4A cells stained with zyxin (green) and Paxillin (red). Boxes mark regions that are shown as higher resolution images.

(C) Quantification of percent of zyxin-positive focal adhesions in control and Flag-Rab40b-4A cells. n ≥ 15 cells per cell line were analyzed. Control - 76.2 ± 2.9 SEM, Flag-Rab40b-4A - 89.7 ± 2.1 SEM.

(D) Western Blot images of cell lysates blotted with anti-FAK, anti-Y397 pFAK or anti-S910 pFAK antibodies. Numbers shown are the average densitometry analysis derived from three independent experiments, relative to tubulin and standardized to control levels. Control::Flag-Rab40b, S910 p = 0.05; Control::Flag-Rab40b-4A, S910 p < 0.005, Y397 p < 0.05.

**Supplemental Figure 3**

(A) Control, Flag-Rab40b and Flag-Rab40-4A cell lines were plated on collagen-coated coverslips, fixed and stained with anti-cortactin (image on left) and anti-CD63 antibodies (image on right). Boxes mark the region that is displayed as higher-resolution image. Arrows point to organelles positive for both, cortactin and CD63. Quantification on the left are the means and SEM.

(B) Western Blot images of Tks5 and p130Cas from control and Flag-Rab40b-4A cells. Numbers represent densitometry analysis from one biological replicate, relative to tubulin and standardized to control levels.

(C) The knock-down efficiency for Rab40b siRNA and Rab40c siRNA as determined by RT-qPCR.

(D) Control, Rab40-3KO-1 and Rab40-3KO-2 cell lines were plated on collagen-coated coverslips, fixed and stained using phalloidin-Alexa594. Boxes mark the region that is displayed as higher-resolution image. Arrows point to stress fibers. Quantification on the right are the means and SEM derived from three independent analyses. Total of 150 cells were analyzed for each condition.

**Supplemental Figure 4**

(A-D) Control, Rab40b-4A and Rab40-3KO cell lines were plated on collagen-coated coverslips, fixed and stained using phalloidin-Alexa594 and anti-non-muscle myosin IIA/B antibodies. Boxes mark the region that is displayed as higher-resolution image. Arrows point to stress fibers. Quantification in D are the means and SEM derived from three independent analyses. Dots represent individual cells analyzed.

(E) Quantification of number of focal adhesions per cell for control, Rab40b-3KO-1 and Rab40b-3KO-2 cells. n ≥ 20 cells per condition.

**Supplemental Figure 5**

(A-B) Control, Flag-Rab40b, Rab40b-4A, and Rab40-3KO cell lines were seeded on collagen-coated coverslips and incubated for 90 minutes. Cells were then fixed and stained using phalloidin-Alexa594. Quantification on the right are the means and SEM derived from three independent analyses. Dots represent individual cells analyzed.

(C) Control, Flag-Rab40b-4A, and Rab40-3KO cell lines were seeded on collagen-coated coverslips. Invadopodia dynamics was then imaged by time-lapse microscopy/DIC. Small line marks the location of quantification shown on the bottom. To visualize the dynamics of the entire lamellipodia see Supplemental Movies 5-8.

**Supplemental Figure 6**

(A) Control or Flag-Rab40b-4A expressing cells were transfected with either GFP-EPLIN-α (green) or GFP-EPLIN-β (green). Cells were then fixed and stained with phalloidin-Alexa594 (red). Line indicates the area analyzed by line scan and shown in (B).

(B-C) EPLIN and actin distribution in lamellipodia was analyzed by line-scan. The location of line scan analysis is shown in (A). Panel see shows the analysis of the distance between actin front at the leading edge and EPLIN front. The data shown are the means and standard error of means derived from 5 different cells.

(E) Representative images of immunohistological stains of EPLIN from tumors grown from each cell line.

**Supplemental Figure 7**

Uncropped Western Blot images shown in previous Figures.

**Supplemental Table 1**

Full results of mass spectrometry experiments

**Supplemental Movie 1**

MDA-MB-231 cells stained with SiRactin (yellow) and DAPI (blue), plated on collagen coated glass bottom dishes and imaged every 10 minutes for 10 hours.

**Supplemental Movie 2**

FLAG-Rab40b-4A expressing cells stained with SiRactin (yellow) and DAPI (blue) plated on collagen coated dishes and imaged every 10 minutes 10 hours.

**Supplemental Movie 3**

MDA-MB-231 cells transfected with GFP-paxillin (green) and stained with SiRactin (red), plated on collagen coated glass bottom dishes and imaged every 4 minutes for 280 minutes.

**Supplemental Movie 4**

FLAG-Rab40b-4A expressing cells transfected with GFP-paxillin (green) and stained with SiRactin (red), plated on collagen coated glass bottom dishes and imaged every 4 minutes for 280 minutes.

**Supplemental Movie 5**

MDA-MB-231 cells were plated on collagen coated glass bottom dishes and imaged by DIC. 50 images with time lapse of 3 seconds were taken for each cell.

**Supplemental Movie 6**

FLAG-Rab40b-4A expressing cells were plated on collagen coated glass bottom dishes and imaged by DIC. 50 images with time lapse of 3 seconds were taken for each cell.

**Supplemental Movie 7**

Rab40-3KO cells were plated on collagen coated glass bottom dishes and imaged by DIC. 50 images with time lapse of 3 seconds were taken for each cell.

