## Supplementary figures and images for "Rab40/Cullin5 complex regulates EPLIN and actin cytoskeleton dynamics during cell migration and invasion"

### Supplemental Figure 1

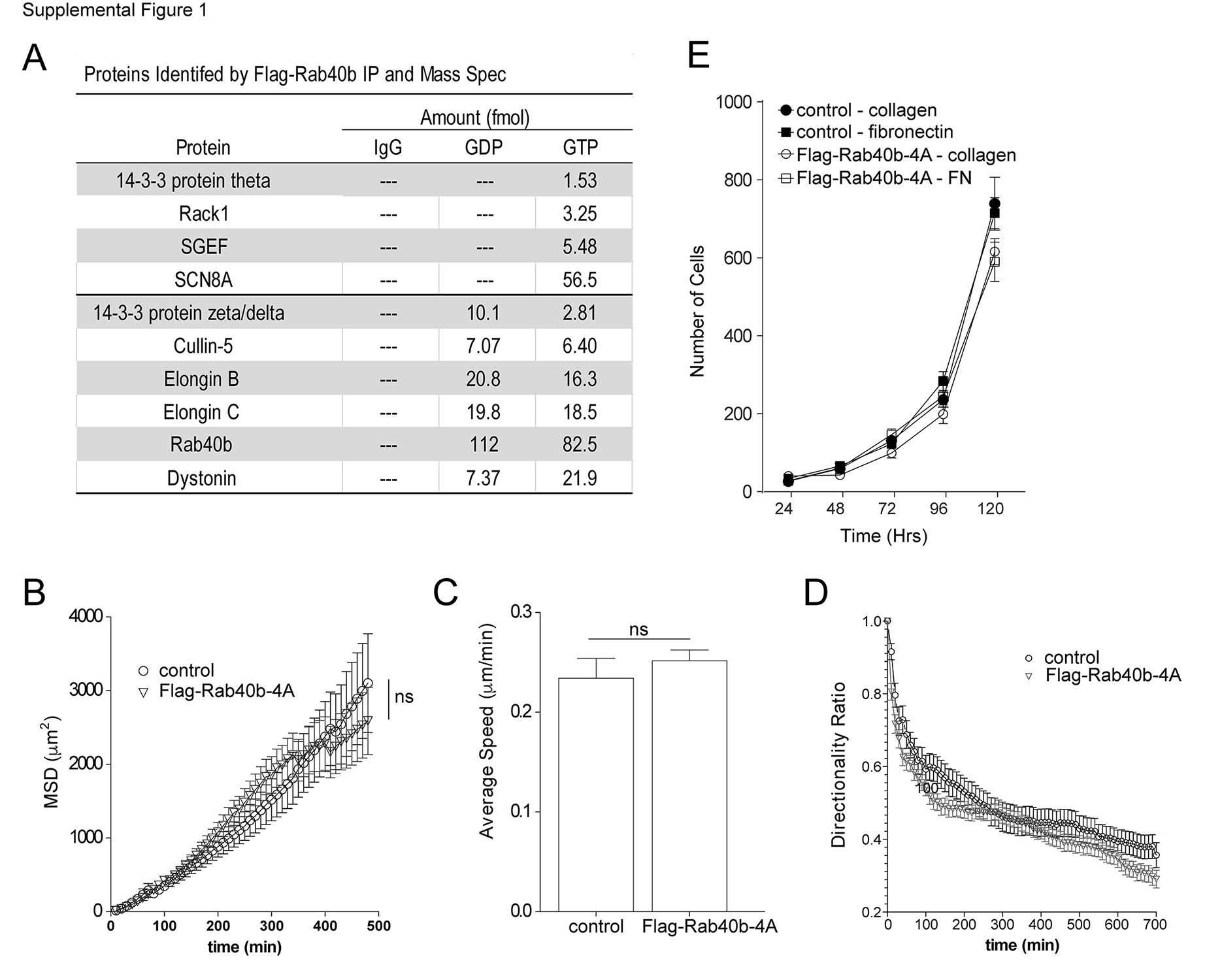

### Supplemental Figure 2

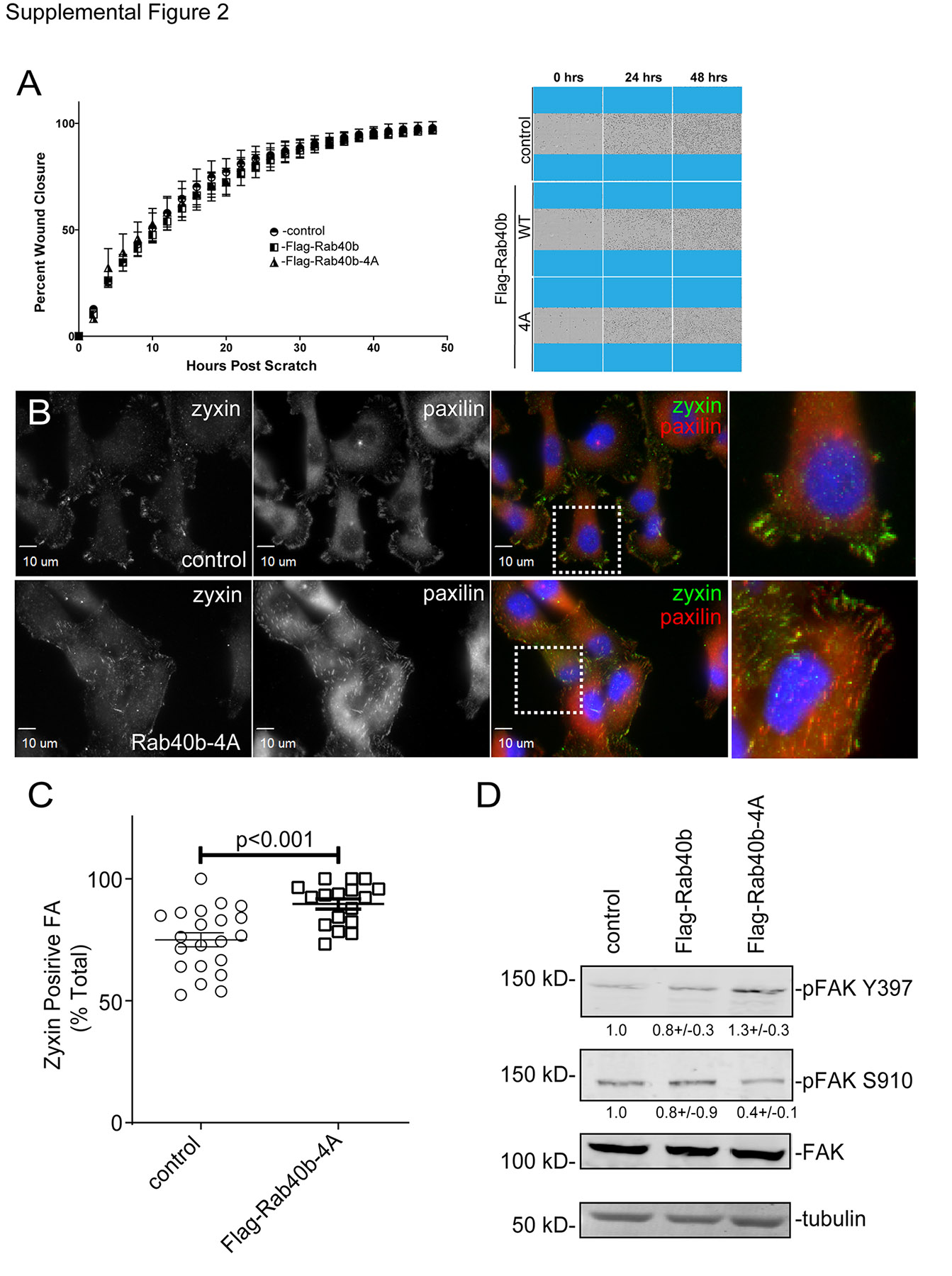

### Supplemental Figure 3

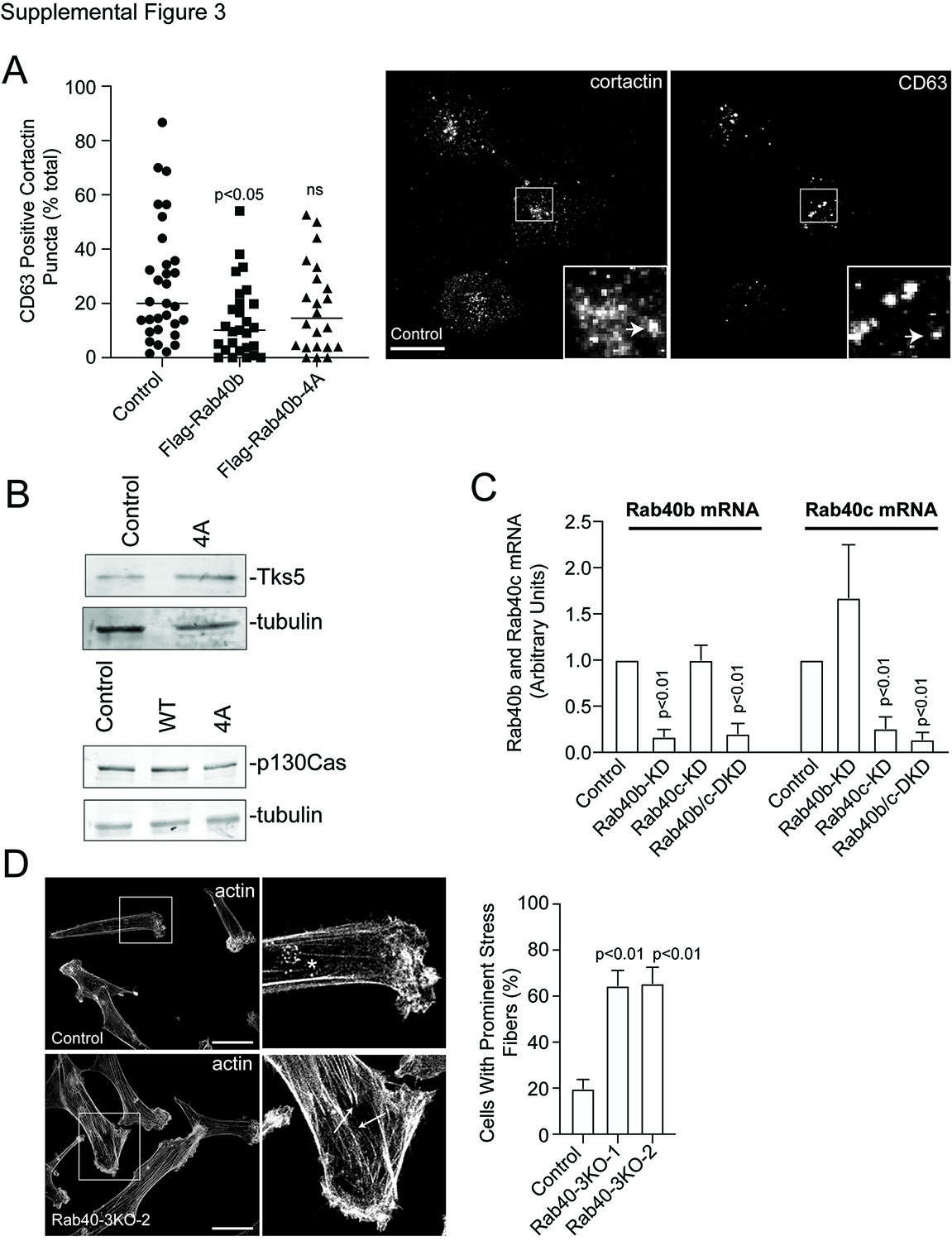

### Supplemental Figure 4

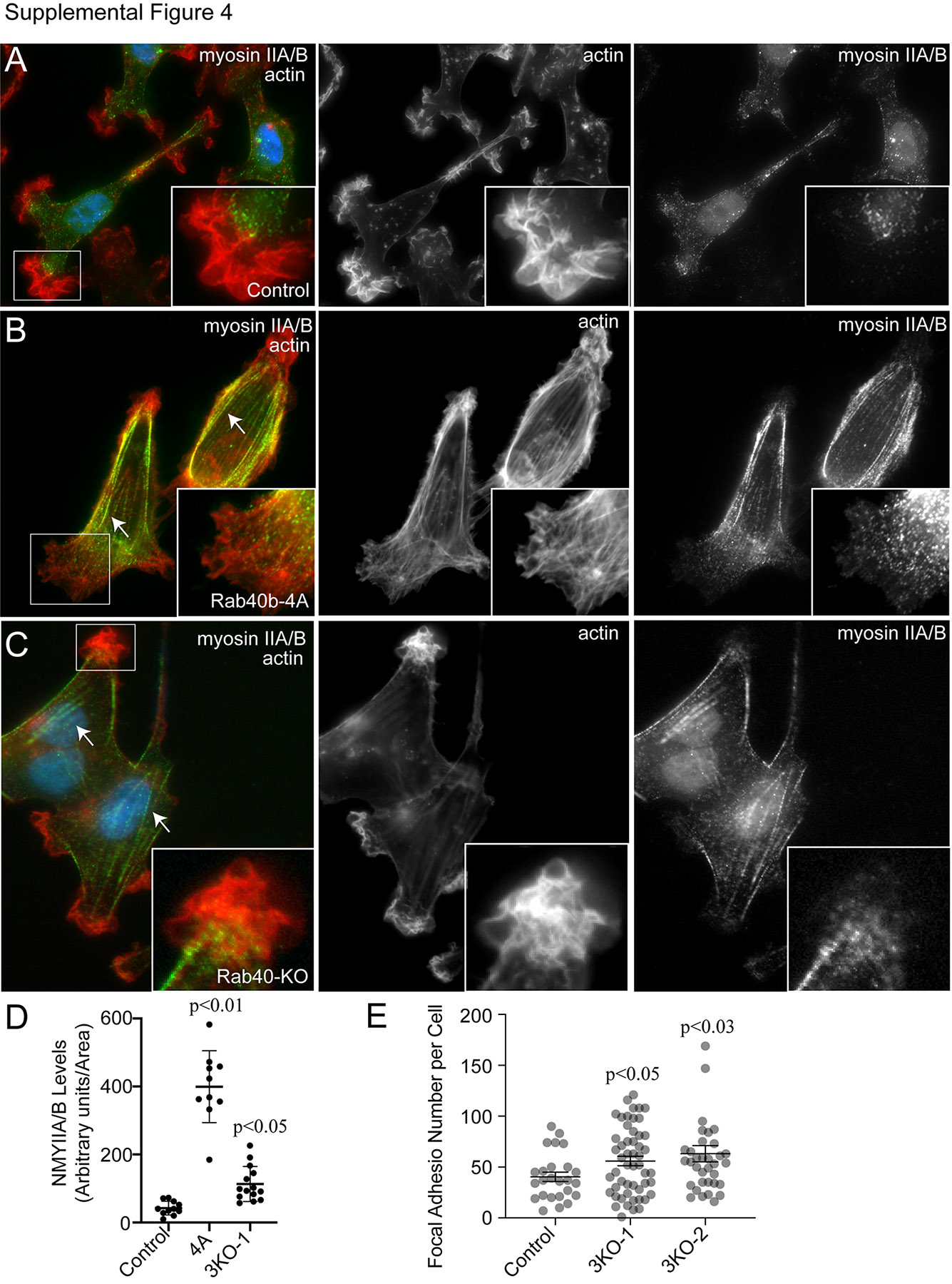

### Supplemental Figure 5

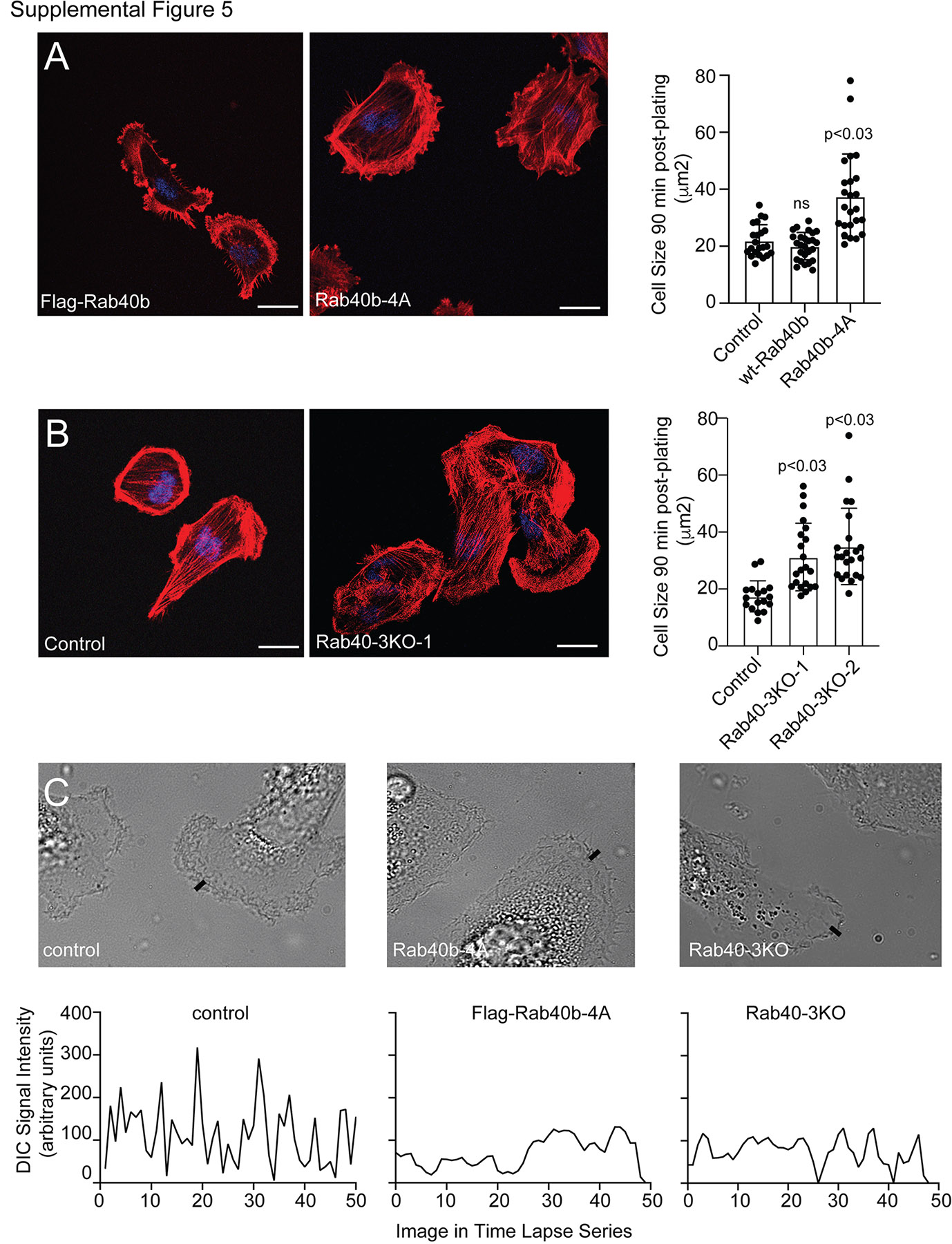

### Supplemental Figure 6

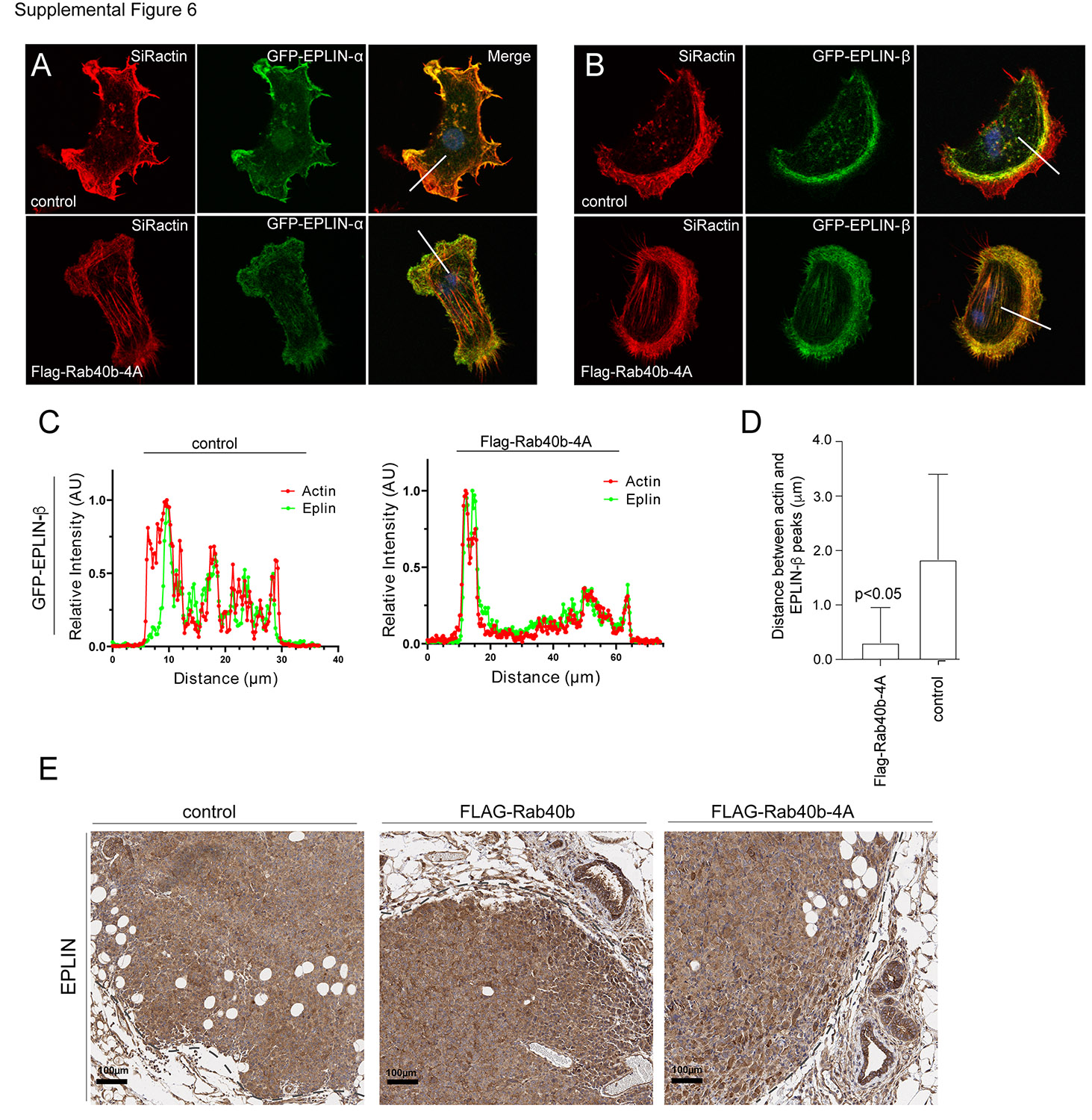
